# Chronic stress induces co-ordinated cortical microcircuit cell type transcriptomic changes consistent with altered information processing

**DOI:** 10.1101/2020.08.18.249995

**Authors:** Dwight F. Newton, Hyunjung Oh, Rammohan Shukla, Keith Misquitta, Corey Fee, Mounira Banasr, Etienne Sibille

**Author notes:** Corresponding Author: Etienne Sibille, PhD, 250 College street, Toronto, ON M5T 1R8, Canada.

## Abstract

Major depressive disorder (MDD) is associated with altered GABAergic and glutamatergic signalling, suggesting altered excitation-inhibition balance (EIB) in cortical mood- and cognition-regulating brain regions. Information processing in cortical microcircuits involves regulation of pyramidal (PYR) cells by Somatostatin-(SST), Parvalbumin-(PV), and Vasoactive intestinal peptide-(VIP) expressing interneurons. Human and rodent studies suggest that impaired PYR-cell dendritic morphology and decreased SST-cell function may mediate altered EIB in MDD. However, knowledge of co-ordinated changes across microcircuit cell types is virtually absent. We thus investigated the co-ordinated transcriptomic effects of UCMS on microcircuit cell types in the medial prefrontal cortex. C57Bl/6 mice, exposed to unpredictable chronic mild stress (UCMS) or control housing for five weeks were assessed for anxiety- and depressive-like behaviours. Microcircuit cell types were laser-microdissected and processed for RNA-sequencing. UCMS-exposed mice displayed predicted elevated behavioural emotionality. Each microcircuit cell type showed a unique transcriptional signature after UCMS. Pre-synaptic functions, oxidative stress response, metabolism, and translational regulation were differentially dysregulated across cell types, whereas nearly all cell types showed down-regulated post-synaptic gene signatures. At the microcircuit level, we observed a shift from distributed transcriptomic co-ordination across cell types in controls towards UCMS-induced increased co-ordination between PYR-, SST- and PV-cells, and a hub-like role for PYR-cells. Lastly, we identified a microcircuit-wide co-expression network enriched in synaptic, bioenergetic, and oxidative stress response genes that correlated with UCMS-induced behaviours. Together, these findings suggest cell-specific deficits, microcircuit-wide synaptic reorganization, and a shift in cortical EIB mediated by increased co-ordinated regulation of PYR-cells by SST- and PV-cells.

## INTRODUCTION

Major depressive disorder (MDD) is a severe psychiatric disorder characterized by low mood, anhedonia, and cognitive dysfunction affecting over 300 million people annually [1, 2]. Conventional monoaminergic antidepressants have poor response rates (50%), resulting in a substantial gap in treatment necessitating novel antidepressant modalities [3]. Moreover, given the neuronal diversity of the brain [4] and the burgeoning understanding of cell-specific pathologies in MDD, a refined approach to understanding its neurobiology and identifying potential antidepressant targets is required [5].

Cortical brain regions, implicated in MDD pathophysiology, are characterized by a repeated and phylogenetically conserved microcircuitry, consisting of glutamatergic pyramidal cells (PYR-cells), regulated by diverse inhibitory, γ-aminobutyric acidergic (GABAergic) interneurons (Figure 1A) [5, 6]. Somatostatin (SST)-expressing cells innervate distal PYR-cell dendrites and regulate corticocortical and thalamocortical inputs [5]. Parvalbumin (PV)-expressing cells innervate the soma and axon hillock of PYR-cells, regulating excitatory output [5]. Vasoactive-intestinal peptide (VIP)-expressing cells innervate SST-cells, disinhibiting PYR-cells [5]. PYR-cell projections onto each interneuron population mediate feedback-inhibition, creating an interconnected structure with an inherent excitation-inhibition balance (EIB). This balance suggests that acute and chronic transcriptional changes within cells are likely co-ordinated across cell types, rather than exclusive to one cell type. We described elsewhere how altered cell type-specific function could affect the EIB, alter information processing and corrupt the cortical microcircuit-mediated neural code [7].

**Figure 1.**
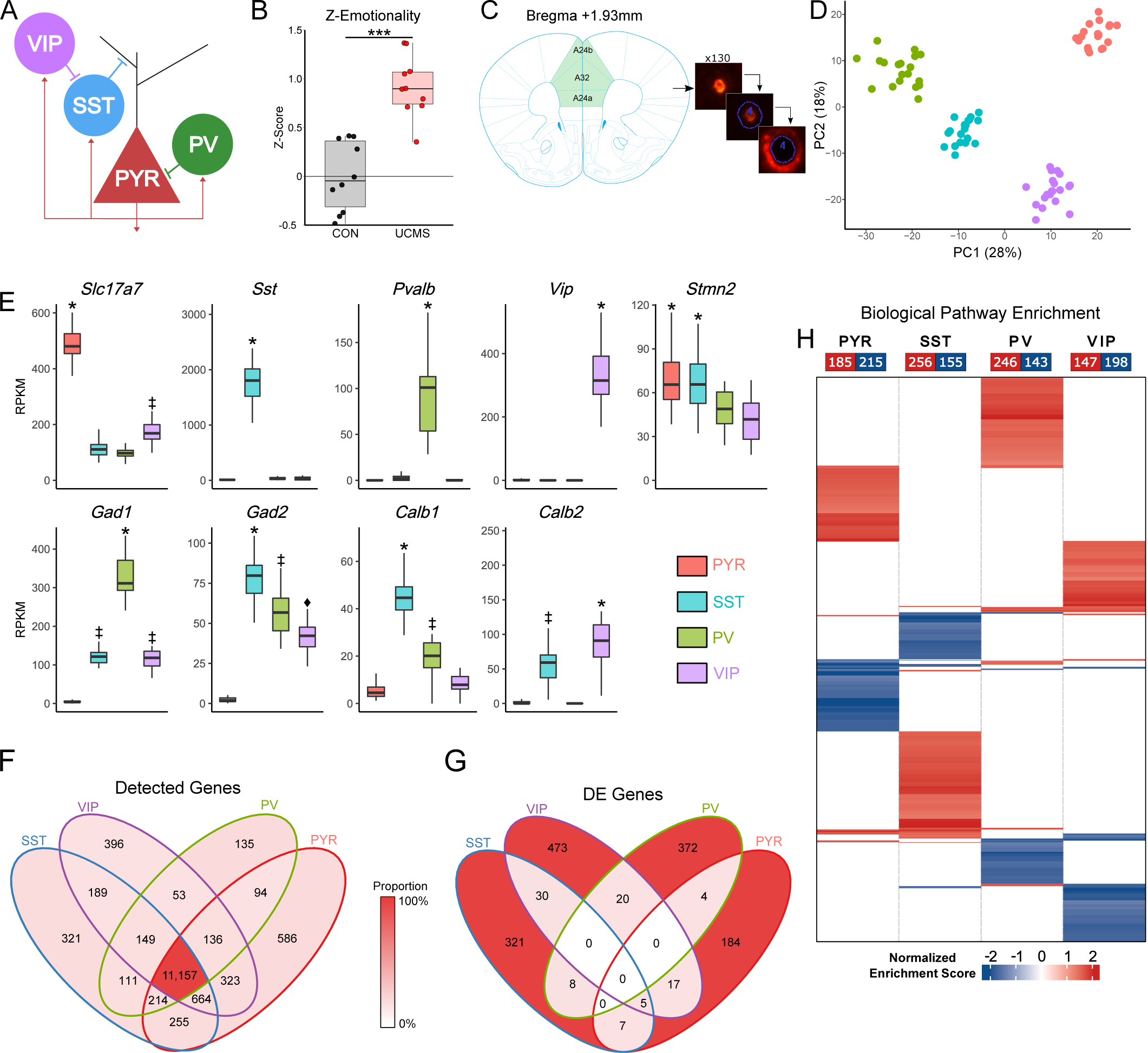
Isolated microcircuit cell types show expected enrichment of molecular markers and cell-specific transcriptomic perturbations. **A)** Illustration of the cortical microcircuit and connectivity patterns of primary cell types. PYR-cells (red) are excitatory glutamatergic cells, regulated by different GABAergic interneurons. Somatostatin-expressing (SST) cells (blue) synapse onto PYR-cell dendrites, providing continuous tonic inhibitory tone. Parvalbumin-expressing (PV) cells (green) synapse onto perisomatic regions and the axon hillock of PYR-cells, and other PV-cells (not shown), to provide phasic inhibition and synchrony of PYR-cell firing across cortical microcircuit columns. Vasoactive-intestinal peptide-expressing (VIP) cells (violet) synapse onto SST-cells, regulating their activity and contributing to the integration of corticocortical and thalamocortical inputs onto PYR-cell dendrites and VIP-cells. PYR-cells form synapses onto each interneuron cell type, mediating feedback inhibition. **B)** Behavioural emotionality Z-scores in control (black) and UCMS-exposed (red) mice. Emotionality is significantly increased (p=7.41×10^−6^) in UCMS-exposed mice. **C)** Delineation of brain regions from which 130 of each microcircuit cell type was extracted. The laser-capture microdissection workflow is visualized, in which a VIP-cell (red fluorescence, image 1) is identified and the laser path is manually traced (blue outline). A space between the cell and laser path prevents the laser from directly damaging cellular contents (image 2). After laser-dissection, a halo surrounding the area previous occupied by the cell results from the reaction of the tissue to the laser (image 3). C) Enrichment of cell type molecular marker expression. *,‡, and ◊ indicate p<1×10^−4^ versus those without the same symbol. **D)** Principle component analysis (PCA) of omnibus gene expression in PYR, SST, PV, and VIP cells. K-means clustering (k=4) confirms that each cell type forms exclusive clusters. **E)** Venn diagram of number of genes detected across cell types. **F)** Venn diagram of number of significant DE (p<0.05, |log_2_ fold-change|>20% genes across cell types. **H)** Heatmap showing significant enriched (p<0.05) biological pathways in each cell type, shaded by normalized enrichment score (blue: down-regulated, red: up-regulated).

The anterior cingulate cortex (ACC) is of particular interest in MDD due to its involvement in mood regulation, showing elevated activity during depressive episodes and normalization with successful pharmacological and non-pharmacological antidepressants [8]. EIB disruptions in MDD manifest *in vivo*, with cortical inhibitory deficits reflecting decreased GABA_A_ and GABA_B_-mediated inhibition, and reduced GABA concentration in cortical brain regions [9, 10]. Decreased expression of GABAergic and glutamatergic synaptic genes were reported in transcriptomic studies of MDD [11-13]. Though technical limitations have limited large-scale gene expression studies to bulk-tissue analyses, masking cell type-specific effects, targeted investigations of cell type-specific molecular markers have repeatedly identified reduced SST expression in the ACC and prefrontal cortex in MDD [14-16]. Importantly, this reduction manifests as reduced expression per-cell, not reduced SST cell density, suggesting impairments in SST-cell function [15].

Further insights into cellular bases of EIB disruptions in MDD have come from mouse stress models, including unpredictable chronic mild stress (UCMS), a paradigm which recapitulates behavioural and neurobiological phenotypes homologous to those observed in MDD [7, 17, 18]. Mice exposed to UCMS show reduced SST expression and increased signalling of the unfolded protein response (UPR) in SST-cells, indicating compromised proteostasis [15, 19]. These deficits suggest impaired SST-mediated inhibition, which is predicted to reduce the filtering of spurious excitatory inputs and decrease the overall signal-to-noise ratio of PYR-cell output, impairing the processing of mood-relevant information. Indeed, mice lacking SST show elevated depressive-like behaviour, suggesting that these deficits may be causally involved in such behaviour [19]. UCMS studies report evidence of PYR-cell deficits, namely reduced spine density, dendritic atrophy, and synapse loss [20-22]. PYR-cells also show reduced expression of post-synaptic glutamatergic markers after UCMS [22, 23]. However, these cellular deficits occur in the context of the cortical microcircuitry, and the concurrent changes occurring in PV and VIP-cells are under-studied [24]. Moreover, the co-ordinated changes occurring across the microcircuitry in either MDD or UCMS, even between SST and PYR-cells, have been inferred from disparate studies and not studied in combination [7, 20, 25].

Stress represents a potential origin for these cellular deficits, given its salience in precipitating depressive episodes and the similarity of UCMS and MDD on multiple investigational levels. Moreover, UCMS provides a model in which to study early pathological changes on the timescale of weeks, versus years of disease burden in MDD [26]. We hypothesized that UCMS induces co-ordinated transcriptomic changes within and across microcircuit cell types and that some of these cellular and molecular changes are related to anxiety and depressive-like behaviours. To test this hypothesis, we used laser-capture microdissection (LCM) to harvest microcircuit cell types from the medial prefrontal cortex (mPFC), roughly homologous to the ACC in humans [27]. RNA sequencing (RNAseq) was employed to obtain cell type-specific, transcriptome-wide, gene expression profiles in mice exposed to UCMS versus controls. We predicted that (1) previously-observed markers of functional and/or structural deficits in SST- and PYR-cells would be replicated, (2) cell types would show unique transcriptional profiles in response to UCMS, and (3) changes would be co-ordinated across cell types, reflecting maintenance of EIB at an altered homeostatic set-point.

## METHODS

See supplementary methods for detailed materials and methods

Twenty male C57Bl/6J mice were exposed to either control or UCMS conditions (n=10/group) for 5-weeks and underwent a battery of anxiety- and anhedonia-like behavioural tests before sacrifice, as previously described [26]. 130 PYR-, SST-, PV-, and VIP-cells/mouse were collected from the mPFC using fluorescent *in situ* hybridization and LCM (Figure 1C) [28]. RNAseq was performed as described [29], using the HiSeq 2500 platform (Illumina, San Diego, CA). Differential expression (DE), gene-set enrichment analysis (GSEA), and weighted gene co-expression analysis (WGCNA) were used to assess transcriptomic changes within and across cell-types [30-32]. DE significance was set at fold-change>20% and p<0.05.

## RESULTS

### UCMS-exposed mice exhibit elevated behavioural emotionality

Mice exposed to 5-weeks of UCMS exhibited an expected increase in behavioural emotionality (p=7.41×10^−6^), an omnibus measure of behaviour test scores (Figure 1B). See all behavioural test results in Supplementary Figure 1.

### RNA-Sequencing reveals robust and microcircuit cell type-specific gene enrichment profiles

RNAseq of LCM-collected single cell types revealed 13 430 genes detected above expression thresholds in PYR-cells, 13 061 in SST-cells, 12 050 in PV-cells, and 13 068 in VIP-cells, with 11 157 expressed in all cell types (Figure 1F). Principal component analysis of gene expression demonstrated robust cell type-specific separation (100% accuracy: k-means clustering, k=4) (Figure 1D) and cell types showed expected enrichment of molecular markers (all p<1×10^−4^) (Figure 1E): *Slc17a7, Sst, Pvalb*, and *Vip* in their respective cell types; *Gad1* (GAD67) and *Gad2* (GAD65) exclusively in interneurons; *Calb1* (calbindin) in SST and PV-cells, and *Calb2* (calretinin) in SST and VIP-cells; *Stmn2*, a pan-neuronal marker, in all cell types.

### UCMS induces microcircuit cell type unique biological changes

DE analysis yielded 217 DE genes (119 up, 98 down) in PYR-cells, 371 (171 up, 200 down) in SST-cells, 406 (168 up, 238 down) in PV-cells, and 545 (269 up, 276 down) in VIP-cells, which were largely unique to each cell type (Figure 1G, gene-lists in Supplementary Table 1). Analysis of altered biological pathways using GSEA identified 400 biological pathways enriched in PYR-cells (185 up, 215 down), 411 in SST-cells (256 up, 155 down), 389 in PV-cells (246 up, 143 down), and 345 in VIP-cells (147 up, 198 down) (Figure 1H). These results were summarized using unbiased clustering of overlapping biological pathways (Figure 2), in parallel with analyzing biological functions with *a priori* links to MDD and UCMS (Supplemental Methods and Supplemental Figure 2). Results are described next for each cell type.

**Figure 2.**
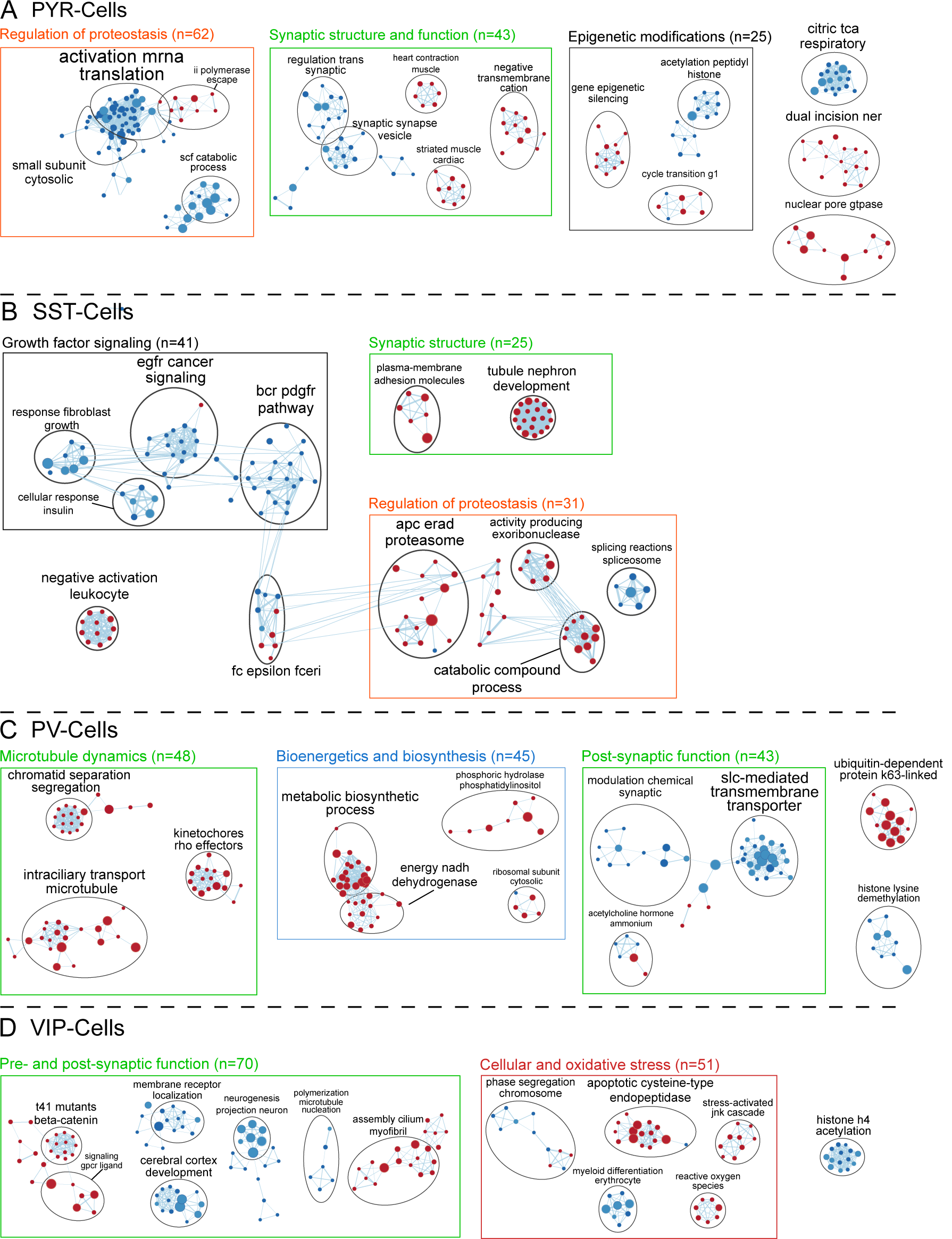
Cluster-based visualization of biological pathways altered by UCMS shows cell-specific profiles of synaptic, bioenergetic, and proteostasis deficits. EnrichmentMap networks displaying GSEA results for (**A**) PYR-cells, (**B**) SST-cells, (**C**) PV-cells, and (**D**) VIP-cells. Nodes indicate gene-sets (pathways), with node size representing the number of genes. Edges indicate the similarity of gene-sets calculated by a 50:50 ratio of the Overlap and Jaccard indices. Red and blue indicate up- and down-regulation, respectively. Clusters defined by Markov clustering were named based on semantic similarity of gene-set names (using the AutoAnnotate Cytoscape package). As cluster names are based on gene-set names, spurious non-neuronal functions can occur (e.g. *tubule nephron development*) and are further characterized by examining shared functions and leading-edge genes. Green boxes indicate clusters related to altered synaptic functions, orange indicates proteostasis and autophagy, blue indicates bioenergetics, red indicates cellular stress, and black indicates functions altered in only one cell type. Bracketed numbers indicate the total number of gene-sets in the group of clusters.

#### PYR-Cells (Figure 2A, Supplementary Table 2)

Three clusters, representing 62 pathways, associated with the regulation of proteostasis, encompassing translational machinery and initiation, proteasome and autophagy, showed down-regulation. These changes were driven by *Eif4h, Psmb6*, and *Bag6*, three genes associated with protein translation and quality control, down-regulated in UCMS and enriched across multiple pathways in each cluster. Five clusters (43 pathways) were associated with synaptic structure and function. These clusters were characterized by down-regulation of neurotransmitter release driven by pre-synaptic (*Snap25, Cplx1*) and post-synaptic (*Grin2d, Nos1*) genes, and up-regulated inhibition of ion transporters and glutamatergic signalling (*Homer1* and *Prcka*). Three clusters (25 pathways) associated with epigenetic modifications were both up- and down-regulated, though no significantly DE genes drove these findings. One cluster (18 pathways) associated with bioenergetics (*citric tca respiratory*) showed down-regulation of oxidative phosphorylation, primarily cytochrome C subunits (*Cox5a, Cox7b, mt-Co3*).

Enrichment of *a priori* biological functions was consistent with cluster-based results, implicating altered synaptic transmission and stress-related pathways (Supplementary Figure 2). Notably, up-regulated *ion channel inhibitor activity* (Normalized Enrichment Score=1.65, p=0.017) pathways may reflect dendrite reorganization processes, driven by *Caprin1* (p=0.0079), a key negative regulator of synaptic plasticity and protein synthesis [33].

These results suggest a general decrease in PYR-cell excitatory function, evidenced by down-regulation of synaptic structure and function genes, compromised proteostasis and reduced mitochondrial function. At the microcircuit-mediated EIB level, these changes suggest reduced PYR-mediated information propagation and feedback activation of local inhibitory cells.

#### SST-Cells (Figure 2B, Supplementary Table 3)

Four clusters (41 pathways) associated with growth factor and neurotrophic signalling pathways, namely insulin, epidermal (EGF), fibroblast (FGF) and platelet-derived growth factors (PDGF) were down-regulated in SST-cells of UCMS-exposed mice. Four clusters (31 pathways) associated with proteostasis, characterized by response to endoplasmic reticulum stress (ER-stress), were up-regulated. These pathways were driven by proteasome, chaperone (*Hspa5* and *Tor1a*), and mRNA degradation genes (*Eri1, Pde5a*, and *Cnot7*). Two clusters (25 pathways) associated with synaptic structure, including post-synaptic cadherin-mediated adhesion (*Smad4, Celsr1, Clstn2*, and *Ctnnd1*) were also up-regulated. Finally, enriched *a priori* functions were consistent with those described above (Supplementary Figure 2), including up-regulated synapse structure, stress-related pathways, cell-adhesion molecules (CAMs; driven by cadherin binding-proteins), and ER-stress (driven by chaperone proteins *Hspa5, Hspa14*, and *Tor1a*).

Overall, these results suggest decreased SST-cell function, evidenced by reduced growth factor and neurotrophic support, altered proteostasis, and increased ER-stress (primarily translational inhibition). At the microcircuit-mediated EIB level, these results suggest decreased SST-cell mediated inhibitory activity of PYR-cells.

#### PV-Cells (Figure 2C, Supplementary Table 4)

Three clusters (48 pathways) associated with microtubule dynamics were up-regulated in PV-cells of UCMS-exposed mice. These clusters were characterized by increased axonal microtubule polymerization, stabilization, and anterograde transport (*Rhot2, Exoc5, Whamm*), and inhibition of retrograde transport along microtubules (*Anapc4*). Four clusters (45 pathways) associated with bioenergetics and biosynthesis were up-regulated, driven by genes involved in glycolysis (*Gpd1*), oxidative phosphorylation (*mt*-*Cytb, mt*-*Atp6*), and nucleotide biosynthesis (*Prps1*). Lastly, 3 clusters (43 pathways) associated with post-synaptic functions were down-regulated, including glutamatergic and cholinergic signalling (*Chrna4, Gria1*), and post-synaptic CAMs (*Dlgap4, Slitrks*), were down-regulated. *A priori* function enrichment was similar to cluster-based results, suggesting up-regulation of ER-stress (*Atf4, Gcn1l1*) and bioenergetics-related pathways, and down-regulation of synaptic-related pathways (Supplementary Figure 2), particularly potassium ion channels (*Kcnk9, Kcnt1, Kcnc3*).

Overall, PV-cell results suggest increased neuronal and cellular bioenergetic activities and decreased glutamatergic and cholinergic signalling input. At the microcircuit-mediated EIB level, these results suggest increased PV-mediated inhibition of PYR-cell output.

#### VIP-Cells (Figure 2D, Supplementary Table 5)

VIP-cells showed bi-directional enrichment in 7 clusters (70 pathways) associated with synaptic structure and function. These clusters suggest complex reorganization of axonal and dendritic cytoskeletons, as each was enriched in pathways and genes involved in both cellular compartments. This included down-regulated neurotrophic factor signalling relevant to cytoskeletal maintenance (*Arsb, Nfasc, Nf2*), and post-synaptic CAMs (*Snap47, Stx1b*), and up-regulated protein phosphatase 2A signalling (*Ppp2re5*), involved in synaptic reorganization. Five clusters (51 pathways) associated with cellular and oxidative stress showed mixed patterns, with decreased regulation and activity of autophagosomes (*Rab11a*), increased mitogen-activated protein kinase (*Map3k5*) signalling, increased oxidative stress response (*Sod2*), and increased ER-stress response (*Bcap31, Hspa9*). *A priori* functions (Supplementary Figure 2) showed consistent down-regulation of synaptic pathways and up-regulation of cellular stress pathways. Down-regulated synaptic structure pathways were reflective of decreased post-synaptic complexes (*Snap47*), axonal growth and function (*Ank1, Nfasc*), and synaptic vesicle release (*Rab3c*). Stress pathways reflected up-regulated response to reactive oxygen species (ROS), driven by *Sod2*, and up-regulated apoptotic signalling through mitogen-activated protein kinase (MAPK) pathways, primarily p38-MAPK signalling (*Map3k5, Dusp10*).

Overall, VIP-cell results suggest a potential atrophy or reorganization of axonal and synaptic compartments, combined with increased oxidative and other cellular stressors. The functional consequences on VIP-cell activity are unclear, though decreased VIP-cell integrity and function are likely. At the microcircuit-mediated EIB level, such changes would contribute to impaired ability of VIP-cells to regulate SST-cell activity, consequently reducing SST-mediated regulation of long-range excitatory inputs onto PYR-cells, consistent with disinhibition of PYR-cells as a primary function of VIP-cells.

### UCMS induces a concerted strengthening of gene co-expression patterns between PYR-cells and SST-/PV-cells

We next used weighted gene co-expression network analysis (WGCNA) to explore putative co-ordinated changes across microcircuitry cell types in response to UCMS, and co-expression signatures of UCMS-induced behaviour (Figure 3A). To this end, we identified gene co-expression modules within individual cell types and determined the degree to which these modules were correlated across cell types. Twenty-six to thirty-three gene co-expression modules were identified per cell type under control or UCMS conditions and summarized by their representative eigengene value. In controls, the eigengene co-expression networks revealed a balanced pattern of correlated modules, suggesting a homogeneous equilibrium across cell types (Figure 3B). In UCMS, a marked shift was observed, with significantly increased module co-expression between PYR and PV-cells (p=1.0×10^−4^), and between PYR and SST-cells (p=0.029), whereas correlations between PYR and VIP-cells did not change (p=0.42).

**Figure 3.**
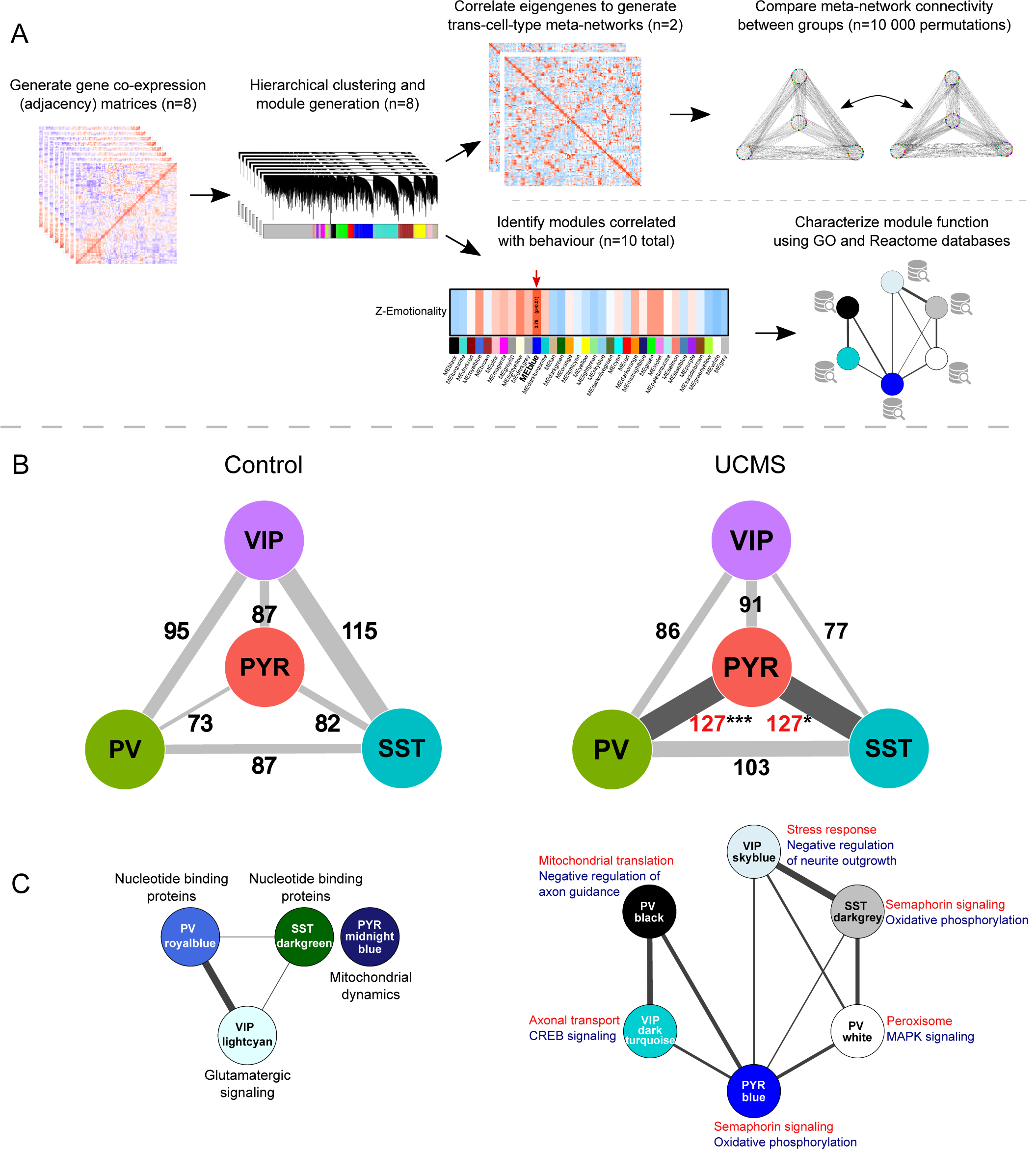
UCMS strengthens transcriptome-wide co-expression between PYR-cells and both SST- and PV-cells. **A)** Analytical workflow employed in co-expression analyses. **B)** Transcriptome-wide co-expression patterns in control (left) and UCMS (right) groups. WGCNA modules generated within cell types were correlated, generating a trans-cell type meta-network. Edges between cell types represent the number of significant pairwise correlations between module eigengenes. Counts in black showed no significant differences between groups, those in red showed a significant increase in co-expression in UCMS. Statistical significance was calculated by permutation tests (n=10 000) of WGCNA results. *p<0.05, ***p<0.001. **C)** Sub-network of WGCNA modules significantly correlated to behaviour in control (left) and UCMS (right) groups. Labels indicate biological functions most significantly enriched in each module using Enrichr. In the UCMS sub-network, functions enriched in up-regulated genes are shown in red, and down-regulated genes in blue.

These results demonstrate an increased correlation of transcriptome-wide changes in SST-, PYR-, and PV-cells following UCMS, suggesting enhanced functional co-ordination. Combined with the cell type-specific functional analyses, these results suggest decreased SST-cell mediated regulation of PYR-cell input, decreased PYR-cell signalling, and increased PV-cell regulation of PYR-cell output, together maintaining the EIB, at a lower degree of activity.

### Gene co-expression modules relevant to UCMS-induced behaviour suggest a co-ordinated response to stress, implicating synaptic reorganization

Finally, to identify co-expression modules and biological functions correlating with UCMS-induced behaviours, we extracted a sub-network of modules with significant eigengene correlations with behavioural emotionality z-scores (Figure 1B). Few modules correlated with behavioural variability in controls (Pearson r=–0.71, 0.64-0.84), which formed a disconnected network (Figure 3C, left). Conversely, a greater, but still small, number of modules were correlated with behaviour after UCMS (r=0.67-0.81). These modules formed an organized sub-network linking the 4 cell types, centered around PYR-cells, as detected by increased PYR-cell hub-related graph measures (Closeness centralization: UCMS=0.629 versus controls=0.208, p=2.86×10^−5^; Betweenness centralization: UCMS=0.320 versus controls=0, p=2.55×10^−4^) (Figure 3C, right). UCMS-associated modules showed enrichment in axonal and dendritic reorganization, bioenergetics, and cellular stress in all cell types (details in Supplementary Table 6).

Taken together, these results implicate PYR-cells as a hub cell type, co-ordinating a microcircuitry-wide response to UCMS, characterized by cellular stress responses, altered metabolism, and synaptic reorganization.

## DISCUSSION

In this study, we investigated cortical microcircuit cell type-specific transcriptomic changes induced by chronic stress (UCMS), circuit-wide co-ordinated adaptations, and transcriptional signatures related to UCMS-induced behavioural deficits. This study also represents the first large-scale transcriptomic analyses of PV-and VIP-cells, in addition to PYR- and SST-cells simultaneously, in UCMS. First, we identified unique transcriptional signatures in each cell type after UCMS. Second, pre-synaptic functions, ROS response, metabolism, and translational regulation were differentially dysregulated across cell types, whereas nearly all cell types showed down-regulation of post-synaptic gene signatures. Third, we observed a shift in cellular coordination, in which UCMS increased co-expression between PYR and both SST and PV-cells. Lastly, we identified a microcircuit-wide co-expression network highly enriched in genes related to synaptic, bioenergetic, and ROS response that correlated with the UCMS-induced behavioural deficits. Together, these findings identify differential effects of UCMS on microcircuit cell types, are consistent with compromised SST- and PYR-cell function [7, 20], identify pathological processes occurring in PV and VIP-cells, and implicate synaptic reorganization processes across the microcircuitry and a potential shift in cortical EIB to a decreased level of activity.

### Microcircuit-wide effects of UCMS

Our cell-specific findings suggest a pattern of synaptic reorganization occurring across the microcircuitry in UCMS versus controls, as illustrated in Figure 4. PYR-cell changes suggest decreased glutamatergic inputs onto all cell types, and SST and VIP-cell changes suggest decreased inhibition of their respective targets. PV-cells on the other hand, are the only cell type which show a transcriptomic profile consistent with increased activity. This may represent a compensatory increase in PV-mediated inhibition of PYR-cell output, following decreased SST-mediated inhibition of PYR-cell inputs, to maintain the EIB of the microcircuitry. These results are strengthened by observations of altered mitochondrial function and ROS response across cell types, internally consistent with the respective increases and decreases in bioenergetic demand. These results are consistent with a substantial body of literature indicating loss of PYR-cell structure and function [34, 35], loss of SST-cell proteostasis and ER-stress [19], and decreased GABAergic and glutamatergic function [25]. Importantly, we identify here for the first time that such deficits co-occur simultaneously.

**Figure 4.**
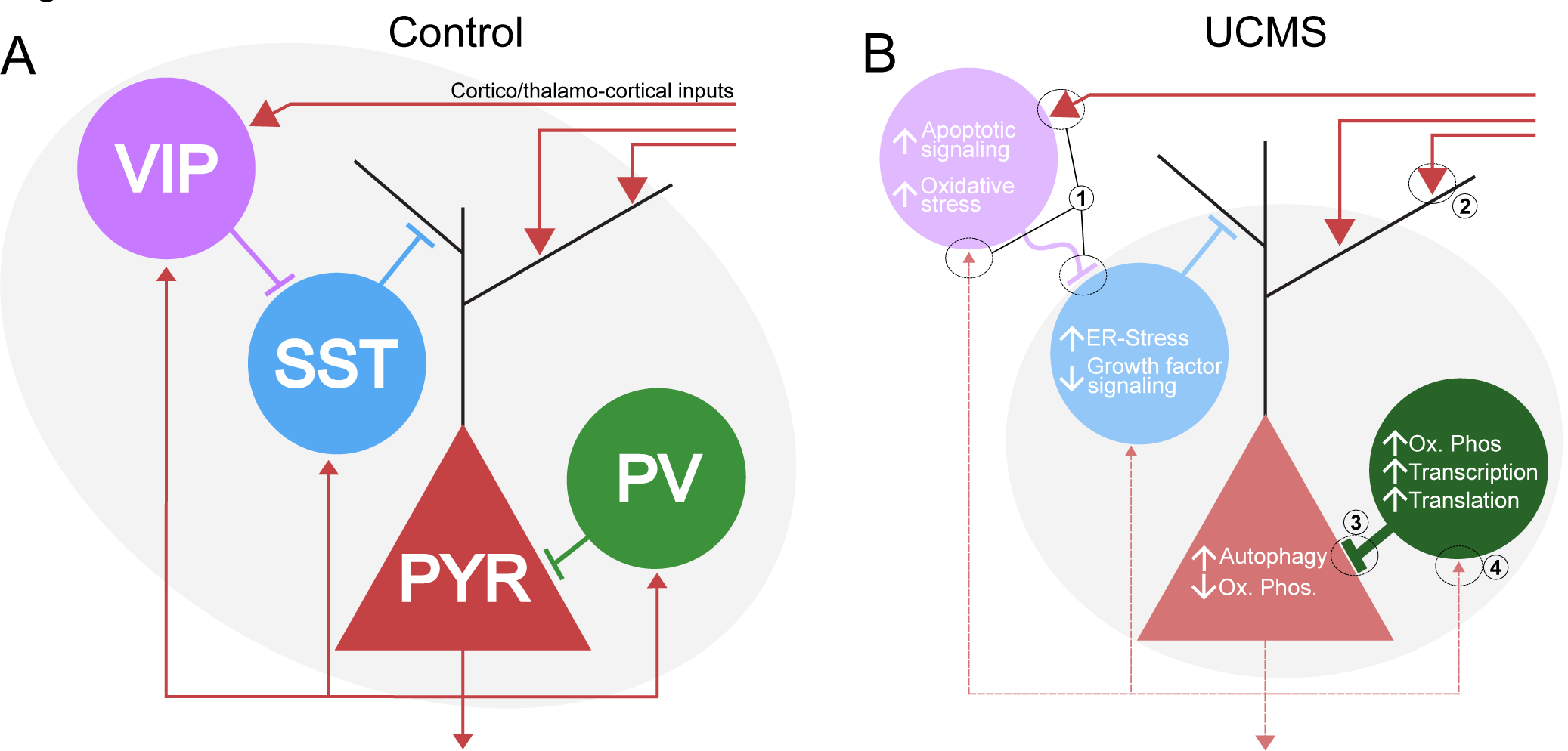
Expected functional consequences of UCMS-induced transcriptomic changes on the cortical microcircuit. **A)** A healthy cortical microcircuit maintains EIB and normal information processing through recruitment and involvement of all cell types (grey outline), where regulation of excitatory input by VIP- and SST-cells is balanced by PV-cell mediated output regulation, ensuring appropriate processing and transfer of PYR-cell mediated excitatory information. **B)** UCMS induces a structural and functional reorganization of the cortical microcircuit. All cell types show perturbations as a result of UCMS, though co-ordinated transcriptomic responses across the microcircuit are more restricted to PYR, PV, and SST-cells (grey outline), suggesting an organized recruitment of these cell types. Intracellular signalling changes, coupled with synaptic changes, suggest decreased activity of PYR, SST, and VIP-cells and increased PV-cells activity: (1) Aberrant axonal and dendritic expression in VIP-cells compromises proper integration of corticocortical and feedback inputs, in addition to impaired transmission of inhibitory signals to SST-cells; (2) Decreased glutamatergic receptor complexes in PYR-cells reduce response to corticocortical and thalamocortical inputs; (3) Increased axonal microtubule and cytoskeletal gene expression, along with increased biosynthesis, suggest increased inhibition of PYR-cells; (4) Decreased post-synaptic glutamatergic complexes in PV-cells suggest decreased feedback signalling by PYR-cells, aside from the general decrease in PYR-cell pre-synaptic gene-expression. These changes may manifest as decreased VIP-/SST-mediated input and increased PV-mediated output regulation of PYR-cells, consequently maintaining the EIB, but potentially negatively impacting the coding and integrity of neural information.

Our co-expression results provided three primary findings. First, UCMS induces a coordinated, potentially compensatory, set of transcriptomic changes occurring primarily between PYR, PV, and SST-cells. This indicates that the integrated response to UCMS is biased towards these cell types, with VIP-cell deficits occurring in relative isolation. Indeed, the emotionality-related subnetwork, showing a convergent enrichment profile of biological functions with cell-specific analyses, suggests that synaptic reorganization, oxidative stress response, and oxidative phosphorylation processes occur in a co-ordinated manner across the microcircuitry. Second, a portion of the cell-specific synaptic, oxidative stress, and bioenergetic disruptions identified in DE analyses are correlated across cell types, in a highly connected manner. This further supports an active, co-ordinated response to UCMS across the microcircuitry, and support our DE results through an independent analytical approach. Third, PYR-cells represented the primary hub cell type in both the meta-network and sub-network analyses. This suggests that PYR-cells are the putative focal point, but not necessarily initiating factor, of the response to UCMS within the microcircuit, and that the PYR-PV-SST complex is a particularly salient target for potential pharmacotherapy.

### Cell type-specific effects of UCMS

We observed a unique transcriptomic profile in each cell type after UCMS. PYR-cells showed deficits in glutamatergic signalling, translation, autophagy, and post-synaptic receptor complexes. PYR-cell deficits in glutamatergic and synaptic gene expression complement our observations of decreased post-synaptic glutamatergic receptors and scaffolding gene expression in PYR, PV, and VIP-cells, which all receive PYR-cell inputs [5]. This suggests pervasive pre-synaptic (and post-synaptic) deficits in PYR-cells across multiple efferent cellular targets, consistent with previous reports of reduced PYR-cell synapses in UCMS and reduced PYR-cell function [35]. Reduced mitochondrial function in PYR-cells is also largely consistent with extant UCMS literature [36, 37]. Autophagy, chaperone, and translational deficits indicate reduced turnover of malfunctioning subcellular components, a process critical for neurons, given their highly dynamic nature [38]. Autophagic machinery is involved in synaptic plasticity and remodelling, particularly the regulation of synaptic vesicles and post-synaptic GABA and NMDA receptors [39]. Autophagic deficits can alter PYR-cell EIB through altered trafficking of post-synaptic receptors, providing internal consistency to these results [40]. Indeed, this may reflect a potential antidepressant modality, given that the rapid-acting antidepressant ketamine increases autophagy [41] and dendritic spine regeneration [42]. Overall, our PYR-cell findings, consistent with existing UCMS literature, suggest reduced PYR-cell activity in UCMS, implicating reduced processing of information in the mPFC and altered regulation and integration of PYR-cell inputs from reduced feedback inhibition.

SST-cells were characterized by reduced growth factor signalling and up-regulation of pre-synaptic CAMs and response to ER-stress after UCMS. ER-stress induces adaptive responses of reduced translation and increased chaperone, ROS generation, protein quality control, and mRNA and protein degradation [43]. SST-cells showed evidence of all such changes, driven by elevated *Hspa5*, a chaperone and primary molecular sensor of misfolded proteins which activates the UPR, one arm of the ER-stress response [43-45]. Notably, this finding replicates previous work from our group showing increased UPR response in SST-cells after UCMS exposure [19]. Reduced EGF, PDGF, and FGF signalling in SST-cells can be interpreted as neurotrophic or cellular identity deficits. These signalling pathways are neurotrophic in the adult CNS, and their inhibition or abolishment has negative cellular and behavioural consequences [46-50]. Though reduced brain-derived neurotrophic factor is implicated in chronic stress and MDD, these other growth factor pathways may represent additional neurotrophic deficits [12, 51-53]. PDGF has protective effects against ROS and NMDA receptor-mediated excitotoxicity [50]. PDGF and FGF are involved in synaptic plasticity and regulation of NMDA receptor activation, and reduction or abolishment of FGF induces anxiety and anhedonia-like behaviour [46-50]. These signalling pathways also contribute to determining GABAergic and glutamatergic cell-fates [50, 54, 55], and reduced function may contribute to loss of cellular identity, similar to natural aging [56]. Overall, SST-cells show reduced cellular integrity, suggestive of a decreased capacity to provide continuous PYR-cell dendritic inhibition. Decreased dendritic inhibitory capability of SST-cells is predicted to reduce gating of corticocortical inputs onto PYR-cells, resulting in increased spontaneous firing and decreased signal-to-noise ratio of PYR-cell output, ultimately altering the integrity and content of this output [7].

PV-cells were characterized by down-regulated post-synaptic receptors, CAMs, and potassium channel expression, and up-regulated mitochondrial function, biosynthetic pathways, and microtubule dynamics. Decreased glutamatergic receptor subunits, scaffolding genes (including *Gria1* and *Dlgap4*), and AMPA-associated K^+^ channels (*Kcnt1*), in addition to decreased post-synaptic CAM expression (*Slitrk* members), suggest reduced excitatory input from PYR-cells, consistent with our observations of pre-synaptic deficits in PYR-cells. AMPA receptors and leak K^+^ channels are critical for the fast-spiking properties of PV-cells which provide phasic inhibition of, and regulate synchronous activity across, PYR-cells [5, 57]. Conversely, polymerization and stabilization of axonal microtubules was increased, suggestive of axonal development. Anterograde organelle transport, a process strongly linked to axonogenesis and synaptogenesis, was increased in PV-cells [58, 59]. Taken together with increased mitochondrial translation and oxidative phosphorylation, this suggests that PV-cells may expand their complex axonal arbors towards PYR and other PV-cells, though the absence of GABA receptor changes in PV-cells suggests PYR-cell specificity [5]. Overall, though PV-cells show deficits in excitatory inputs, synaptic changes suggest increased cellular activity and pre-synaptic function. Implications for the microcircuitry would be greater, but less regulated and temporally constrained (i.e. decreased phasic and increased tonic), inhibition of PYR-cell output. This increased, but less co-ordinated, control of PYR-cell output may exacerbate the impact of reduced SST-cell inhibition through further dyssynchrony of PYR-cells and loss of signal-to-noise ratio.

Lastly, VIP-cells appeared to show compromised pre- and post-synaptic structure and function, in addition to decreased cellular integrity and neurotrophic signalling. In both enrichment analyses, VIP-cells showed deficits in post-synaptic markers, namely decreased CAMs and dendritic structure genes. These findings are consistent with pre-synaptic deficits in PYR-cells discussed above. Down-regulation of genes involved in axonal structure and vesicle release would likewise suggest a decreased inhibition of SST-cells. These findings may reflect adaptations of VIP-cells to reduced PYR and SST-cell function, though they may stem from increased intracellular apoptotic signalling, indicative of cellular stress. *Map3k5* (also *Ask1*), an initiator of apoptosis, showed up-regulation along with downstream effects including increased oxidative stress response and reduced autophagosome activity [60]. Increased apoptotic signalling is also observed in bulk-tissue sequencing of the mPFC in UCMS [37, 61]. Despite VIP-cells constituting only 3-4% of the cortex, apoptotic signalling is low in control conditions and may be detectable in bulk-tissue if robustly increased in a low-abundance cell type [5]. Lastly, VIP-cell densities were unchanged (Supplementary Figure 3), precluding actual cell death, though *Sod2* was up-regulated and has been shown to forestall apoptosis [62]. Increased apoptotic signalling and oxidative stress in VIP-cells may contribute to an overall decrease in cell function, consistent with previous report of reduced VIP peptide expression in UCMS [24], and contribute to the loss of maintenance of synaptic functions, perhaps leading to impaired regulation of SST-cells.

### Overall predicted implications for the cortical microcircuit

Overall, our DE and co-expression results are suggestive of a shift in microcircuit EIB towards a lower level of overall activity (Figure 4). This reduction is characterized by reduced PYR-cell activity indicating a lower level of information processing in the mPFC, reduced SST-cell function suggesting decreased input regulation of PYR-cells, and increased PV-cell function suggesting increased output control. VIP-cells show synaptic deficits unco-ordinated with the other cell types (most importantly SST-cells), and as such appear to play a decreased role in PYR-cell regulation, albeit this role is primarily via modulation of SST-cells which show deficits outlined above. Reduced input regulation and tighter control of excitatory outputs may result in a less flexible microcircuit that is unable to filter spurious and noisy inputs and able to propagate a more narrow bandwidth of signals. Though these changes appear to maintain the EIB of the microcircuit, their consequences would be a corruption of information output to other cortical and subcortical brain regions.

### Limitations and Future studies

This study is not without limitations. First, only male mice were used, precluding conclusions about effects in females. Depression disproportionately affects females versus males at an approximate ratio of 2:1, and some pathological features, namely reduced SST, are more robust in females than males [16]. Second, mice received 5-weeks of UCMS, thus these transcriptomic changes represent both stress-induced cellular responses and compensatory adaptations of other cell types. Our data is cross-sectional and cannot assess the sequence of cellular deficits. Third, few of the DE or GSEA enrichment results passed correction for multiple comparisons, and as such these findings should be considered exploratory. Fourth, our sample size was moderate (n=10/group), though comparable to other mouse RNAseq studies [37, 61, 63, 64]. Lastly, the cell types investigated in this study are not fully unique, and further divisions of SST- and PV-cells for instance can be made [4, 65].

Future studies should focus on replicating our cell type-specific and microcircuit-wide findings, and investigate the time-course of these changes, to determine if such findings are independent and concomitant or potentially sequential in nature, and ultimately determine their potential as antidepressant modalities. As we found that cell types respond uniquely to UCMS, future studies should also determine the cell type-specific effects of existing and novel ADs on transcriptional profiles, particularly those with synaptic effects such as ketamine. Lastly, studies utilizing post-mortem MDD brain samples should determine if MDD-associated cell specific changes parallel UCMS-induced changes observed here.

## ACKNOWLEDGEMENTS

We thank the Centre for Addiction and Mental Health Sequencing Facility for assistance running the Illumina HiSeq2500 platform and Gary Bader for input on co-expression analyses. Co-expression computations were performed on the CAMH Specialized Computing Cluster (SCC). The SCC is funded by the Canada Foundation for Innovation, Research Hospital Fund. We thank Netta Ussyshkin for help with cell counting.

This work was supported by a Canadian Institute of Health Research Project Scheme Grant (PJT-153175). D.N. was supported by funding provided by the Ontario Graduate Scholarship and Brain Canada, in partnership with Health Canada, for the Canadian Open Neuroscience Platform initiative. C.F. was supported by a CAMH Discovery Fund fellowship and Ontario Graduate Scholarship during the studies. M.B. is supported by a NARSAD young investigator award from the Brain & Behaviour Research Foundation (#24034) and the CAMH Discovery Seed Fund and the Canadian Institutes of Health Research (PJT-165852). This work was also supported by the Campbell Family Mental Health Research Institute.

## REFERENCES

1. Hasin DS, Sarvet AL, Meyers JL, et al. Epidemiology of adult dsm-5 major depressive disorder and its specifiers in the united states. JAMA Psychiatry 2018; 75(4): 336–346.

2. American Psychiatric Association. Diagnostic and Statistical Manual of Mental Disorders, 5th Edition: Washington, DC, 2013.

3. Prins J, Olivier B, Korte SM. Triple reuptake inhibitors for treating subtypes of major depressive disorder: the monoamine hypothesis revisited. Expert opinion on investigational drugs 2011; 20(8): 1107–1130.

4. Tasic B, Yao Z, Graybuck LT, Smith KA, Nguyen TN, Bertagnolli D et al. Shared and distinct transcriptomic cell types across neocortical areas. Nature 2018; 563(7729): 72–78.

5. Tremblay R, Lee S, Rudy B. GABAergic Interneurons in the Neocortex: From Cellular Properties to Circuits. Neuron 2016; 91(2): 260–292.

6. Northoff G, Sibille E. Cortical GABA neurons and self-focus in depression: a model linking cellular, biochemical and neural network findings. Molecular psychiatry 2014; 19(9): 959.

7. Fee C, Banasr M, Sibille E. Somatostatin-positive GABA Interneuron Deficits in Depression: Cortical Microcircuit and Therapeutic Perspectives. Biological psychiatry 2017.

8. Agid Y, Buzsaki G, Diamond DM, Frackowiak R, Giedd J, Girault J-A et al. How can drug discovery for psychiatric disorders be improved? Nature reviews Drug discovery 2007; 6(3): 189–201.

9. Levinson AJ, Fitzgerald PB, Favalli G, Blumberger DM, Daigle M, Daskalakis ZJ. Evidence of Cortical Inhibitory Deficits in Major Depressive Disorder. Biological psychiatry 2010; 67(5): 458–464.

10. Schur RR, Draisma LW, Wijnen JP, Boks MP, Koevoets MG, Joels M et al. Brain GABA levels across psychiatric disorders: A systematic literature review and meta-analysis of (1) H-MRS studies. Human brain mapping 2016; 37(9): 3337–3352.

11. Choudary PV, Molnar M, Evans SJ, Tomita H, Li JZ, Vawter MP et al. Altered cortical glutamatergic and GABAergic signal transmission with glial involvement in depression. Proceedings of the National Academy of Sciences of the United States of America 2005; 102(43): 15653–15658.

12. Chang L-C, Jamain S, Lin C-W, Rujescu D, Tseng GC, Sibille E. A Conserved BDNF, Glutamate- and GABA-Enriched Gene Module Related to Human Depression Identified by Coexpression Meta-Analysis and DNA Variant Genome-Wide Association Studies. PloS one 2014; 9(3): e90980.

13. Kang HJ, Adams DH, Simen A, Simen BB, Rajkowska G, Stockmeier CA et al. Gene expression profiling in postmortem prefrontal cortex of major depressive disorder. The Journal of neuroscience : the official journal of the Society for Neuroscience 2007; 27(48): 13329–13340.

14. Douillard-Guilloux G, Lewis D, Seney ML, Sibille E. Decrease in somatostatin-positive cell density in the amygdala of females with major depression. Depression and anxiety 2017; 34(1): 68–78.

15. Seney ML, Tripp A, McCune S, Lewis DA, Sibille E. Laminar and cellular analyses of reduced somatostatin gene expression in the subgenual anterior cingulate cortex in major depression. Neurobiology of disease 2015; 73: 213–219.

16. Tripp A, Kota RS, Lewis DA, Sibille E. Reduced somatostatin in subgenual anterior cingulate cortex in major depression. Neurobiology of disease 2011; 42(1): 116–124.

17. Piantadosi SC, French BJ, Poe MM, Timic T, Markovic BD, Pabba M et al. Sex-Dependent Anti-Stress Effect of an α5 Subunit Containing GABAA Receptor Positive Allosteric Modulator. Frontiers in pharmacology 2016; 7: 446–446.

18. Sun HL, Zhou ZQ, Zhang GF, Yang C, Wang XM, Shen JC et al. Role of hippocampal p11 in the sustained antidepressant effect of ketamine in the chronic unpredictable mild stress model. Translational psychiatry 2016; 6: e741.

19. Lin LC, Sibille E. Somatostatin, neuronal vulnerability and behavioural emotionality. Molecular psychiatry 2015; 20(3): 377–387.

20. Duman RS, Sanacora G, Krystal JH. Altered Connectivity in Depression: GABA and Glutamate Neurotransmitter Deficits and Reversal by Novel Treatments. Neuron 2019; 102(1): 75–90.

21. Kang HJ, Voleti B, Hajszan T, Rajkowska G, Stockmeier CA, Licznerski P et al. Decreased expression of synapse-related genes and loss of synapses in major depressive disorder. Nature medicine 2012; 18(9): 1413–1417.

22. Li N, Liu R-J, Dwyer JM, Banasr M, Lee B, Son H et al. Glutamate N-methyl-D-aspartate Receptor Antagonists Rapidly Reverse Behavioural and Synaptic Deficits Caused by Chronic Stress Exposure. Biological psychiatry 2011; 69(8): 754–761.

23. Duman RS, Aghajanian GK, Sanacora G, Krystal JH. Synaptic plasticity and depression: new insights from stress and rapid-acting antidepressants. Nature medicine 2016; 22(3): 238.

24. Banasr M, Lepack A, Fee C, Duric V, Maldonado-Aviles J, DiLeone R et al. Characterization of GABAergic marker expression in the chronic unpredictable stress model of depression. Chronic stress (Thousand Oaks, Calif) 2017; 1.

25. Northoff G, Sibille E. Why are cortical GABA neurons relevant to internal focus in depression? A cross-level model linking cellular, biochemical and neural network findings. Molecular psychiatry 2014; 19(9): 966–977.

26. Nikolova YS, Misquitta KA, Rocco BR, Prevot TD, Knodt AR, Ellegood J et al. Shifting priorities: highly conserved behavioural and brain network adaptations to chronic stress across species. Translational psychiatry 2018; 8(1): 26.

27. George Paxinos KF. The Mouse Brain in Stereotaxic Coordinates. 4th Edition edn. Academic Press 2012.

28. Rocco BR, Oh H, Shukla R, Mechawar N, Sibille E. Fluorescence-based cell-specific detection for laser-capture microdissection in human brain. Scientific Reports 2017; 7(1): 14213.

29. Shukla R, Prevot TD, French L, Isserlin R, Rocco BR, Banasr M et al. The Relative Contributions of Cell-Dependent Cortical Microcircuit Aging to Cognition and Anxiety. Biological psychiatry 2019; 85(3): 257–267.

30. Langfelder P, Horvath S. WGCNA: an R package for weighted correlation network analysis. BMC bioinformatics 2008; 9: 559.

31. Love MI, Huber W, Anders S. Moderated estimation of fold change and dispersion for RNA-seq data with DESeq2. Genome biology 2014; 15(12): 550.

32. Subramanian A, Tamayo P, Mootha VK, Mukherjee S, Ebert BL, Gillette MA et al. Gene set enrichment analysis: a knowledge-based approach for interpreting genome-wide expression profiles. Proc Natl Acad Sci U S A 2005; 102(43): 15545–15550.

33. Solomon S, Xu Y, Wang B, David MD, Schubert P, Kennedy D et al. Distinct structural features of caprin-1 mediate its interaction with G3BP-1 and its induction of phosphorylation of eukaryotic translation initiation factor 2alpha, entry to cytoplasmic stress granules, and selective interaction with a subset of mRNAs. Molecular and cellular biology 2007; 27(6): 2324–2342.

34. Duman RS, Aghajanian GK, Sanacora G, Krystal JH. Synaptic plasticity and depression: new insights from stress and rapid-acting antidepressants. Nature medicine 2016; 22(3): 238–249.

35. McEwen BS, Bowles NP, Gray JD, Hill MN, Hunter RG, Karatsoreos IN et al. Mechanisms of stress in the brain. Nature neuroscience 2015; 18(10): 1353.

36. Bansal Y, Kuhad A. Mitochondrial Dysfunction in Depression. Curr Neuropharmacol 2016; 14(6): 610–618.

37. Tordera RM, Garcia-García AL, Elizalde N, Segura V, Aso E, Venzala E et al. Chronic stress and impaired glutamate function elicit a depressive-like phenotype and common changes in gene expression in the mouse frontal cortex. European Neuropsychopharmacology 2011; 21(1): 23–32.

38. Lee J-A. Neuronal autophagy: a housekeeper or a fighter in neuronal cell survival? Exp Neurobiol 2012; 21(1): 1–8.

39. Tomoda T, Yang K, Sawa A. Neuronal Autophagy in Synaptic Functions and Psychiatric Disorders. Biological psychiatry 2019.

40. Sumitomo A, Yukitake H, Hirai K, Horike K, Ueta K, Chung Y et al. Ulk2 controls cortical excitatory-inhibitory balance via autophagic regulation of p62 and GABAA receptor trafficking in pyramidal neurons, vol. 27 2018.

41. Gassen NC, Rein T. Is There a Role of Autophagy in Depression and Antidepressant Action? Frontiers in Psychiatry 2019; 10(337).

42. Moda-Sava RN, Murdock MH. Sustained rescue of prefrontal circuit dysfunction by antidepressant-induced spine formation. 2019; 364(6436).

43. Hetz C. The unfolded protein response: controlling cell fate decisions under ER stress and beyond. Nature Reviews Molecular Cell Biology 2012; 13: 89.

44. Bertolotti A, Zhang Y, Hendershot LM, Harding HP, Ron D. Dynamic interaction of BiP and ER stress transducers in the unfolded-protein response. Nature cell biology 2000; 2(6): 326.

45. Lee AS. The ER chaperone and signaling regulator GRP78/BiP as a monitor of endoplasmic reticulum stress. Methods (San Diego, Calif) 2005; 35(4): 373–381.

46. Yamanaka Y, Kitano A, Takao K, Prasansuklab A, Mushiroda T, Yamazaki K et al. Inactivation of fibroblast growth factor binding protein 3 causes anxiety-related behaviours. Molecular and Cellular Neuroscience 2011; 46(1): 200–212.

47. Birey F, Kloc M, Chavali M, Hussein I, Wilson M, Christoffel Daniel J et al. Genetic and Stress-Induced Loss of NG2 Glia Triggers Emergence of Depressive-like Behaviours through Reduced Secretion of FGF2. Neuron 2015; 88(5): 941–956.

48. Brooks LR, Enix CL, Rich SC, Magno JA, Lowry CA, Tsai P-S. Fibroblast growth factor deficiencies impact anxiety-like behaviour and the serotonergic system. Behavioural Brain Research 2014; 264: 74–81.

49. Di Re J, Wadsworth PA, Laezza F. Intracellular Fibroblast Growth Factor 14: Emerging Risk Factor for Brain Disorders. Front Cell Neurosci 2017; 11: 103.

50. Funa K, Sasahara M. The roles of PDGF in development and during neurogenesis in the normal and diseased nervous system. J Neuroimmune Pharmacol 2014; 9(2): 168–181.

51. Duman RS. Neurobiology of stress, depression, and rapid acting antidepressants: remodeling synaptic connections. Depression and anxiety 2014; 31(4): 291–296.

52. Oh H, Piantadosi SC, Rocco BR, Lewis DA, Watkins SC, Sibille E. The Role of Dendritic Brain-Derived Neurotrophic Factor Transcripts on Altered Inhibitory Circuitry in Depression. Biological psychiatry 2019; 85(6): 517–526.

53. Tripp A, Oh H, Guilloux JP, Martinowich K, Lewis DA, Sibille E. Brain-derived neurotrophic factor signaling and subgenual anterior cingulate cortex dysfunction in major depressive disorder. The American journal of psychiatry 2012; 169(11): 1194–1202.

54. Garcez RC, Teixeira BL, Schmitt Sdos S, Alvarez-Silva M, Trentin AG. Epidermal growth factor (EGF) promotes the in vitro differentiation of neural crest cells to neurons and melanocytes. Cellular and molecular neurobiology 2009; 29(8): 1087–1091.

55. Stevens HE, Smith KM, Maragnoli ME, Fagel D, Borok E, Shanabrough M et al. Fgfr2 is required for the development of the medial prefrontal cortex and its connections with limbic circuits. The Journal of neuroscience : the official journal of the Society for Neuroscience 2010; 30(16): 5590–5602.

56. Goh JO. Functional Dedifferentiation and Altered Connectivity in Older Adults: Neural Accounts of Cognitive Aging. Aging and disease 2011; 2(1): 30–48.

57. Goldberg EM, Jeong H-Y, Kruglikov I, Tremblay R, Lazarenko RM, Rudy B. Rapid Developmental Maturation of Neocortical FS Cell Intrinsic Excitability. Cerebral Cortex 2010; 21(3): 666–682.

58. Mattson MP, Gleichmann M, Cheng A. Mitochondria in neuroplasticity and neurological disorders. Neuron 2008; 60(5): 748–766.

59. Chang DT, Reynolds IJ. Differences in mitochondrial movement and morphology in young and mature primary cortical neurons in culture. Neuroscience 2006; 141(2): 727–736.

60. Hatai T, Matsuzawa A, Inoshita S, Mochida Y, Kuroda T, Sakamaki K et al. Execution of Apoptosis Signal-regulating Kinase 1 (ASK1)-induced Apoptosis by the Mitochondria-dependent Caspase Activation. Journal of Biological Chemistry 2000; 275(34): 26576–26581.

61. Liu Y, Yang N, Zuo P. cDNA microarray analysis of gene expression in the cerebral cortex and hippocampus of BALB/c mice subjected to chronic mild stress. Cellular and molecular neurobiology 2010; 30(7): 1035–1047.

62. Kannan K, Jain SK. Oxidative stress and apoptosis. Pathophysiology : the official journal of the International Society for Pathophysiology 2000; 7(3): 153–163.

63. Laine MA, Trontti K, Misiewicz Z, Sokolowska E, Kulesskaya N, Heikkinen A et al. Genetic Control of Myelin Plasticity after Chronic Psychosocial Stress. eneuro 2018; 5(4): ENEURO.0166-0118.2018.

64. Manners MT, Yohn NL, Lahens NF, Grant GR, Bartolomei MS, Blendy JA. Transgenerational inheritance of chronic adolescent stress: Effects of stress response and the amygdala transcriptome. Genes, Brain and Behaviour 2019; 18(7): e12493.

65. Tasic B, Menon V, Nguyen TN. Adult mouse cortical cell taxonomy revealed by single cell transcriptomics. 2016; 19(2): 335–346.

