## Supplementary Methods for "Chronic stress induces co-ordinated cortical microcircuit cell type transcriptomic changes consistent with altered information processing"

**Table of Contents**

**Supplementary Methods:**

- ***Experimental Animals***
- ***UCMS Protocol***
- ***Behavioural Testing***
- ***Statistical Analysis of Behavioural Tests***
- ***Sample Collection, Library Preparation, and RNA-Sequencing***
- ***Cell Type-Specificity Assessment***
- ***Differential Expression (DE) and Biological Pathway Enrichment Analysis***
- ***Co-Expression Analyses***
- ***Cell Densitometry***

**Supplementary Figures:**

- **Supplementary Figure 1.** UCMS-exposed mice show elevated behavioural emotionality.
- **Supplementary Figure 2.** Overall enrichment profiles and analysis of a priori biological functions across cell-types.
- **Supplementary Figure 3.** Density of cortical microcircuit cell types in control and UCMS groups.

**Supplementary Tables:**

- **Supplementary Table 1.** Differential expression results in each cell type.
- **Supplementary Table 2.** Leading-edge analysis of EnrichmentMap-derived pathway clusters in PYR-cells.
- **Supplementary Table 3.** Leading-edge analysis of EnrichmentMap-derived pathway clusters in SST-cells.
- **Supplementary Table 4.** Leading-edge analysis of EnrichmentMap-derived pathway clusters in PV-cells.
- **Supplementary Table 5.** Leading-edge analysis of EnrichmentMap-derived pathway clusters in VIP-cells.
- **Supplementary Table 6.** Enrichr analysis of WGCNA modules correlated with emotionality scores in control and UCMS groups.

**Methods**

***Experimental Animals***

All experiments were performed with male C57Bl/6J mice (Jackson Laboratory, Saint Constant, Canada) exposed to UCMS (n=10) or control housing (n=10). Controls were group-housed with littermates whereas UCMS-exposed mice were single-housed. Housing conditions consisted of a 12/12-hour light-dark cycle and constant room temperature (20-23°C). All mice were provided with food and water *ad libitum*, except during UCMS stressors or when required for behavioural testing. All procedures were approved by the Centre for Addiction and Mental Health Animal Care Committee and are in accordance with the Canadian Animal Care Committee.

***UCMS Protocol***

UCMS consisted of a variable sequence of mild social-environmental stressors occurring 2-4 times daily for 5 weeks. Stressors included forced bath (~2cm of water in cage for 15 min), light cycle disruptions or reversal, social stress (cage change with other mice), predator odor (exposure to bobcat or fox urine for 20 min – 1 hour), wet bedding, tilted cage (45° angle), and restraint (50ml conical tube with air hole for 15-30 min), reduced space, or bedding/nestlet removal. UCMS schedule was performed as previously described [1]. Control mice were handled regularly by the behavioural experimenter (KM). Mice were sacrificed 12 hours after the last stressor and 48 hours after the last behavioural test and brains were frozen with dry ice and stored at -80^o^C.

***Behavioural Testing***

Behavioural testing occurred during week 4 and 5, during which UCMS was continued with 1-2 daily stressors to maintain the effects of UCMS during testing. Behavioural tests including elevated plus maze (EPM), open-field (OF), novelty-suppressed feeding (NSF), novelty-induced hypophagia (NIH), sucrose consumption test (SCT), phenotyper test (PT), and a general locomotor activity test.

*Coat State/Fur Quality*

Coat state of 7 anatomical landmarks of the mouse body (head, neck, dorsal coat, ventral coat, forepaws, hindpaws, and tail) was assessed each week, by the same experimenter blinded to group assignment. The fur quality at each landmark was given a score of 0, 0.5, or 1 from maintained to unkempt.

*EPM*

Mice were placed individually into the centre of an elevated (55cm) and dimly illuminated plus maze. The plus maze consisted of two closed arms with walls and two open arms, both opposite each other. The exploration of mice was tracked for 10 minutes, recorded from an overhead video camera. Mouse movement was analyzed using AnyMaze software (Wood Dale, IL, USA), determining the time spent in, and number of entries into, the open and closed arms. Decreased time and entries into the open arms indicated increased anxiety-like behaviour.

*OF*

Mice were placed individually into the centre of a dimly illuminated open field (70cm x 70cm, x 33cm height). Movement within the chamber was tracked from an overhead camera for 10 minutes and analyzed with AnyMaze software. A zone comprising the centre third of the field (23cm x 23cm) was delineated and the time spent, and number of entries, in the zone was determined. A decreased amount of time and number of entries into this centre zone indicated increased anxiety-like behaviour.

*NSF*

Mice were food deprived overnight for 16 hours, after which they were placed on one side of a novel, dimly illuminated arena (clean mouse cage). Four food pellets were placed in the middle of the arena, and the latency to approach the food and latency to bite were both recorded (in seconds). The test was terminated upon the mouse biting the food and lasted a maximum of 12 minutes. Afterwards, mice were returned to their home cages where the experimental paradigm was repeated. Greater latency to approach and bite in a novel environment was indicative of increased anxiety-like behaviour

*NIH*

This experimental paradigm involved a 2-day period in which mice are habituated to a sweetened-condensed milk solution. Habituation consisted of placing a Petri dish containing 1ml of a 33% sweetened-condensed milk solution in the home cage of each mouse for 1 hour. The day following habituation, mice were placed on one end of the home cage, and the milk solution was placed on the opposite end of the cage. The home cage latency to drink was measured during the 5-minute test. The day following home-cage assessment, the paradigm was repeated in a novel cage for a maximum of 10 minutes. A decreased time to approach and drink the solution in a novel environment indicated increased anxiety-like behaviour, with an anhedonia-like behavioural component.

*SCT*

Mice were first habituated to a 1% sucrose solution for 48 hours (in which 1% sucrose solution was available *ad libitum*), after which mice were fluid deprived overnight (16 hours). The next day, the 1% sucrose solution was returned for 1-hour and the amount of sucrose solution consumed was measured. After the test, tap water was made available *ad libitum*. Two days later, mice were deprived of water overnight (16 hours), after which the same 1-hour consumption test was performed with tap water. A decreased consumption of sucrose, when water consumption remained unchanged, indicated increased anhedonia-like behaviour.

*PT*

The phenotyper test is a novel test adapted by our research group that assesses home cage-like behaviour and light-induced neophobia in response to a 1-hour spotlight challenge, over a 14-hour measurement period[2]. Mice were placed in PhenoTyper apparatuses (Noldus, Leesburg, VA, USA) containing an overhead infrared-sensitive camera tracking movement in three pre-defined zones (shelter, food, and water-containing zones). A baseline behavioural measurement was performed over the first four testing hours (7:00pm – 11:00pm), corresponding to the first four dark-cycle hours. After the baseline reading, a white LED spotlight was illuminated for 1-hour, centred over the food zone. Time spent in each zone is measured each hour of the test using the Noldus Ethovision software package. We calculated “residual avoidance” as a measure of anxiety-like behaviour, indicating the persistent avoidance of the food-zone in the hour after the light-challenge had been removed with respect to the amount of time spent during the light-challenge. Residual avoidance is calculated as follows:

$$Residual avoidance=100\times\frac{M-i}{M}$$

Where:

$M$= Mean difference in food-zone time spent in the hour after the light challenge v.s. the hour of the light challenge, among all control mice

$i$=Individual mouse difference in food-zone time spent in the hour after the light challenge v.s. the hour of the light challenge

*Locomotor Activity*

Mice were placed in standard housing cages for 20 minutes in a dimly illuminated room and tracked using an overhead-mounted video camera. Total distance travelled (measured using AnyMaze) was used as a measurement of gross locomotor activity.

***Statistical Analysis of Behavioural Tests***

Behavioural tests were summarized using behavioural test Z-scoring, in which test measures are expressed in terms of the number of standard deviations they lie from the mean of the control group. The following measures (Table A) were Z-scored, then averaged by test to create a test-specific Z-score[3].

| **Table A.** Behavioural measures used for Z-scores. | |
| --- | --- |
| **Test** | **Measure** |
| EPM | Time in open arms  Entries into open arm (% of total) |
| OF | Time in centre zone  Entries into centre zone  Distance travelled in centre zone |
| NSF | Latency to bite (novel cage) |
| NIH | Latency to drink (novel cage) |
| SPT | Sucrose consumed |
| PT | Residual avoidance (food zone) |

These Z-scores were then averaged for an omnibus “behavioural emotionality” Z-score.

Coat state was analyzed using repeated-measures ANOVA to assess effects of stress, time, and stress x time interaction. Post-hoc t-tests by week were performed with a Bonferroni correction. All other behavioural tests were compared between groups using independent-samples t-tests. Significance was defined as p<0.05. Statistical and bioinformatic analyses were performed in R version 3.5.1[4].

***Sample Collection, Library Preparation, and RNA-Sequencing***

12µm coronal sections at cingulate areas 24a, 24b, and 32 (bregma: +1.53 – 1.93) representing the mPFC (Figure 2A), were cut at -20°C using a CM1950 Cryostat (Leica Microsystems, Wetzlar, Germany). Sections were thaw-mounted onto RNase-free polyethylene-naphthalate slides (Leica Microsystems, Wetzlar, Germany) for laser-capture microdissection (LCM) applications. Additional sections were mounted onto RNase-free glass slides for cell density measurements.

A rapid fluorescent *in situ* hybridization (FISH) protocol was used to stain sections with cDNA probes (Advanced Cell Diagnostics, Newark, CA) specific to *Slc17a7* (vesicular glutamate transporter 1), *Sst*, *Pvalb*, or *Vip* to visualize CM cell types, as previously described[5, 6]. 130 cells of each cell type were collected from each mouse using an LMD7 laser microdissection microscope (Leica Microsystems, Wetzlar, Germany). RNA was extracted using a PicoPure RNA isolation kit (Thermo Fisher Scientific, Waltham, MA), and libraries were prepared using the SMARTer Stranded Total RNA-Seq Kit v2-Pico Input Mammalian (Clontech, Mountain View, CA). Library fragment size distribution was determined using a Bioanalyzer (Agilent Technologies, Santa Clara, CA), and sequencing was performed on an Illumina HiSeq 2500 sequencer (Illumina, San Diego, CA) to generate 2x125bp paired-end reads. Reverse reads showed lower and more variable Q30 scores and were thus excluded. Forward reads were trimmed (5’:9bp, 3’:21bp) to remove regions with the highest variation in read quality. Reads passing quality control were aligned to the GRCm38 mouse reference genome ([ftp.ensembl.org/pub/release-86/fasta/mus_musculus/dna/](ftp://ftp.ensembl.org/pub/release-86/fasta/mus_musculus/dna/)) using HiSat2[7] and GenomicAlignments[8] .

***Cell Type-Specificity Assessment***

| **Table B.** Cell type-specific markers | |
| --- | --- |
| Gene | Expected cell type(s) showing specific expression |
| *Slc17a7* | PYR-cells |
| *Sst* | SST-cells |
| *Pvalb* | PV-cells |
| *Vip* | VIP-cells |
| *Gad1* | All interneurons |
| *Gad2* | All interneurons |
| *Calb1* | SST and PV-cells |
| *Calb2* | SST and VIP-cells |
| *Stmn2* | All cell types |

Cell type-specificity was assessed by determining relative expression of known molecular markers and data-driven methods. **Table B** outlines the known cell type markers used [9], and the expected cell types in which they are expressed. Expression of each marker gene of interest was expressed in reads per kilobase of transcript per million mapped reads (RPKM) and compared across cell types using one-way ANOVA with a Bonferroni correction of post-hoc independent-samples t-tests.

Data-driven cell type-specificity was assessed by principal component analysis (PCA). PCA was performed on the top 500 most variable genes across all samples, and samples were plotted by the first and second components (accounting for 46% of the variance in our data). K-means clustering (k=4, 10,000 iterations, 10,000 starting points) was performed, and the known cell type identity of each sample was used as an external validation measure to determine percentage accuracy.

***Differential Expression (DE) and Biological Pathway Enrichment Analysis***

After alignment of reads to exons, DE analysis was performed using DESeq2[10]. To account for differences in cell-specific presence or expression levels of genes, we filtered genes before DE analysis on a cell type-wise basis. In each cell type, genes were excluded if they had less than 10 reads and greater than 2/3 of samples with a 0 count. Contrasts for the effect of UCMS in each cell type were examined. As few genes survived transcriptome-wide multiple-comparison correction[11], exploratory statistical significance was set at p<0.05 and log_2_ fold-change (LFC) >20% and further analyses were performed at the biological pathway level.

Gene Set Enrichment Analysis (GSEA)[12] was performed using Wald statistic-ranked gene lists, EnrichmentMap gene-set database[13], and default GSEA parameters, permuted 10,000 times. An unbiased, hypothesis-generating enrichment analysis of GSEA results within cell types was performed using Cytoscape[13]. Significantly enriched gene-sets were visualized based on degree of shared genes, determined by a 50:50 ratio of the Jaccard and Overlap coefficients. The Jaccard and Overlap coefficients are calculated as follows:

$Jaccard=\frac{Intersection of sets}{Union of sets}$	$Overlap=\frac{Intersection of sets}{Smaller of the two sets}$

The Overlap coefficient provides a more appropriate measure of similarity when comparing sets which differ greatly in size (occurring when GSEA results are restricted to gene sets of 15-500 genes, whereas the Jaccard is more appropriate for gene-sets of similar size. Equally weighting the two is one method of capturing similarity amongst all pairwise comparisons of gene-sets with a single metric[13]. The resulting networks of gene-sets were then grouped using graph-based Markov clustering. Clusters were grouped and characterized by semantic similarity of gene-set names[14] and known functions of leading-edge genes (i.e. those driving the enrichment signal). Leading-edge genes shared amongst at least 5 gene-sets in each cluster, and which passed significance thresholds in DE analysis, were used in this characterization.

Enrichment of biological functions with *a priori* links to MDD and UCMS was determined by exploring parent-child relationships, as previously described[6]. The biological functions were chosen based on literatures reviews of MDD and UCMS neurobiology, and all biological functions selected have been implicated in pathological processes[15-19]. Briefly, Gene Ontology (GO) terms reflective of each of the following processes disrupted in MDD/UCMS were selected (Table C): general synaptic transmission, synaptic structure, GABAergic signalling, glutamatergic signalling, inflammatory response, neurotrophins, cellular and oxidative stress, bioenergetics, and monoaminergic signalling. The number of enriched nested (child) terms within the selected GO terms (parents) was determined in a cell type-specific manner. The distribution of normalized enrichment scores (NES) across all cell types and relative enrichment levels of *a priori* functions is shown in Supplementary Figure 2. The GO-term lists are not exhaustive but represent a common theme to be compared across cell types in a consistent manner.

| **Table C**. GO terms included in *a priori* enrichment analysis (parent-child associations). | | | | | | |
| --- | --- | --- | --- | --- | --- | --- |
| **Biological Function** | **GO ID** | **GO Term** |  | **Biological Function** | **GO ID** | **GO Term** |
| General Neurotransmission | GO:0007268 | chemical synaptic transmission |  | Inflammation | GO:0006955 | immune response |
|  | GO:1902847 | regulation of neuronal signal transduction |  |  | GO:0002437 | inflammatory response to antigenic stimulus |
|  | GO:0098960 | postsynaptic neurotransmitter receptor activity |  |  | GO:0050776 | regulation of immune response |
|  | GO:0016247 | channel regulator activity |  |  | GO:0005125 | cytokine activity |
|  | GO:0038023 | signaling receptor activity |  |  | GO:0004896 | cytokine receptor activity |
|  | GO:0030594 | neurotransmitter receptor activity |  |  | GO:0006954 | inflammatory response |
|  | GO:0005326 | neurotransmitter transporter activity |  | Neurotrophins | GO:0031548 | regulation of brain-derived neurotrophic factor receptor signaling pathway |
|  | GO:0015075 | ion transmembrane transporter activity |  |  | GO:0060398 | regulation of growth hormone receptor signaling pathway |
|  | GO:0030182 | neuron differentiation |  |  | GO:1990089 | response to nerve growth factor |
| Synaptic Structure | GO:0098794 | postsynapse |  |  | GO:1990090 | cellular response to nerve growth factor stimulus |
|  | GO:0098793 | presynapse |  |  | GO:1990792 | cellular response to glial cell derived neurotrophic factor |
|  | GO:0045202 | synapse |  |  | GO:0005030 | neurotrophin receptor activity |
|  | GO:0097458 | neuron part |  | Cellular/Oxidative Stress | GO:0033554 | cellular response to stress |
|  | GO:0097060 | synaptic membrane |  |  | GO:0071500 | cellular response to nitrosative stress |
|  | GO:0050808 | synapse organization |  |  | GO:0034599 | cellular response to oxidative stress |
|  | GO:0007416 | synapse assembly |  |  | GO:0080135 | regulation of cellular response to stress |
|  | GO:0007155 | cell adhesion |  |  | GO:0016491 | oxidoreductase activity |
| GABA | GO:0007214 | gamma-aminobutyric acid signaling pathway |  |  | GO:0055114 | oxidation-reduction process |
|  | GO:0051932 | synaptic transmission, GABAergic |  |  | GO:0036473 | cell death in response to oxidative stress |
|  | GO:0060080 | inhibitory postsynaptic potential |  |  | GO:0006950 | response to stress |
|  | GO:0097154 | GABAergic neuron differentiation |  | Bioenergetics | GO:0006091 | generation of precursor metabolites and energy |
|  | GO:0009450 | gamma-aminobutyric acid catabolic process |  |  | GO:0044262 | cellular carbohydrate metabolic process |
| Glutamate | GO:0035249 | synaptic transmission, glutamatergic |  |  | GO:0005739 | mitochondrion |
|  | GO:1900449 | regulation of glutamate receptor signaling pathway |  |  | GO:0007210 | serotonin receptor signaling pathway |
|  | GO:0007216 | G-protein coupled glutamate receptor signaling pathway |  |  | GO:0099589 | serotonin receptor activity |
|  | GO:0060079 | excitatory postsynaptic potential |  |  | GO:1901338 | catecholamine binding |
|  | GO:0008066 | glutamate receptor activity |  |  | GO:0051937 | catecholamine transport |
|  | GO:1905962 | glutamatergic neuron differentiation |  |  | GO:1901615 | organic hydroxy compound metabolic process |
|  | | |  |  | GO:0003357 | noradrenergic neuron differentiation |
|  |  |  |  |  | GO:0071867 | response to monoamine |

***Co-Expression Analyses***

Weighted gene co-expression network analysis (WGCNA)[20] was used to interrogate co-ordinated gene expression patterns occurring within and across cell types. WGCNA was performed using the union of al genes expressed in each cell type (n=14,391 genes), in control and UCMS groups (n=8 networks). Additional pre-analytical filtering was performed using the “goodSamplesGenes” function from the WGCNA package. Scale-free topology (R^2^<0.9) was achieved at variable soft-thresholding β levels for each cell type-specific network, and a β of 18 was used for all networks for consistency of comparison and mean network connectivity across networks – as per WGCNA guidelines (<https://horvath.genetics.ucla.edu/html/CoexpressionNetwork/Rpackages/WGCNA/faq.html>). Module eigengene meta-networks were used to relate gene-expression across cell types in controls and UCMS groups separately. Meta-networks were generated from Pearson correlation matrices of all eigengenes, excluding intra-cellular correlations, hard-thresholded to include only significant Pearson correlations (p<0.05).

Significance of changes in meta-network connectivity between control and UCMS groups were determined using permutation testing (n=10,000). In each permutation, subjects were randomly assigned to “Control” and “UCMS” groups, and the WGCNA workflow was performed from the initial co-expression matrix step. The number of significant connections between cell types in the meta-networks, corrected for the number of modules in each cell type, was determined. This was used to calculate empirical p-values for the effect of UCMS on the transcriptomic connectivity between cell types.

Separate analyses were performed on a subset of the above WGCNA meta-networks, containing only modules with significant (p<0.05) eigengene correlations with behavioural emotionality Z-scores. These modules were visualized with Cytoscape, and the biological functions represented by each module was characterized using Enrichr and adjusted Fisher’s exact tests[21]. GO and Reactome databases were used in this enrichment analysis, and up/down regulated gene lists were analyzed separately. Closeness and betweenness centralization were determined for each network (UCMS and control) and compared between groups using a permuted jackknife-based approach. In this “leave *n* out” approach, similar to cross-validation, we calculated closeness and betweenness centralization in each group for all possible combinations of subjects in which two were randomly removed. The resulting measures were used to estimate the variance of each measure for each experimental group, and comparisons were performed using independent-samples t-tests. A permutation-based approach as above, where the WGCNA networks were re-generated for each permutation, was not feasible for this analysis as the genes comprising each module of the emotionality-correlated subnetwork would not be consistent and thus the connectivity not reflective of the same coordination in biological processes.

***Cell Densitometry***

Double-label FISH of mPFC sections targeting *Slc17a7* and *Sst,* or *Pvalb* and *Vip,* counterstained with DAPI, was performed to determine changes in cell type density. RNAscope staining was performed according to the manufacturer’s instructions. Six random sites of the mPFC were imaged using an Olympus IX3 confocal microscope (Olympus, Tokyo, Japan). Images were deconvolved using the AutoQuant Blind algorithm in Slidebook version 6.0 (Intelligent Imaging Innovations, Denver, CO) to reduce background, and intensities were normalized across images. Cells containing 10 or more cell-marker grains, and overlapping with DAPI, were manually quantified by a single experimenter. Cell type densities were compared across groups using independent-samples t-tests.

References

1.	Nikolova YS, Misquitta KA, Rocco BR, Prevot TD, Knodt AR, Ellegood J *et al.* Shifting priorities: highly conserved behavioural and brain network adaptations to chronic stress across species. *Translational psychiatry* 2018; **8**(1)**:** 26.

2.	Prevot TD, Misquitta KA, Fee C, Newton DF, Chatterjee D, Nikolova YS *et al.* Residual avoidance: A new, consistent and repeatable readout of chronic stress-induced conflict anxiety reversible by antidepressant treatment. *Neuropharmacology* 2019; **153:** 98-110.

3.	Guilloux JP, Seney M, Edgar N, Sibille E. Integrated behavioural z-scoring increases the sensitivity and reliability of behavioural phenotyping in mice: relevance to emotionality and sex. *Journal of neuroscience methods* 2011; **197**(1)**:** 21-31.

4.	Team RC. R: A Language and Environment for Statistical Computing. Vienna, Austria, 2018.

5.	Rocco BR, Oh H, Shukla R, Mechawar N, Sibille E. Fluorescence-based cell-specific detection for laser-capture microdissection in human brain. *Scientific Reports* 2017; **7**(1)**:** 14213.

6.	Shukla R, Prevot TD, French L, Isserlin R, Rocco BR, Banasr M *et al.* The Relative Contributions of Cell-Dependent Cortical Microcircuit Aging to Cognition and Anxiety. *Biological psychiatry* 2019; **85**(3)**:** 257-267.

7.	Kim D, Langmead B, Salzberg SL. HISAT: a fast spliced aligner with low memory requirements. *Nature Methods* 2015; **12:** 357.

8.	Lawrence M, Huber W, Pagès H, Aboyoun P, Carlson M, Gentleman R *et al.* Software for Computing and Annotating Genomic Ranges. *PLOS Computational Biology* 2013; **9**(8)**:** e1003118.

9.	Tremblay R, Lee S, Rudy B. GABAergic Interneurons in the Neocortex: From Cellular Properties to Circuits. *Neuron* 2016; **91**(2)**:** 260-292.

10.	Love MI, Huber W, Anders S. Moderated estimation of fold change and dispersion for RNA-seq data with DESeq2. *Genome biology* 2014; **15**(12)**:** 550.

11.	Bourgon R, Gentleman R, Huber W. Independent filtering increases detection power for high-throughput experiments. *Proceedings of the National Academy of Sciences* 2010; **107**(21)**:** 9546-9551.

12.	Subramanian A, Tamayo P, Mootha VK, Mukherjee S, Ebert BL, Gillette MA *et al.* Gene set enrichment analysis: A knowledge-based approach for interpreting genome-wide expression profiles. *Proceedings of the National Academy of Sciences* 2005; **102**(43)**:** 15545-15550.

13.	Merico D, Isserlin R, Stueker O, Emili A, Bader GD. Enrichment Map: A Network-Based Method for Gene-Set Enrichment Visualization and Interpretation. *PloS one* 2010; **5**(11)**:** e13984.

14.	Kucera M, Isserlin R, Arkhangorodsky A, Bader G. AutoAnnotate: A Cytoscape app for summarizing networks with semantic annotations [version 1; peer review: 2 approved]. *F1000Research* 2016; **5**(1717).

15.	Boku S, Nakagawa S, Toda H, Hishimoto A. Neural basis of major depressive disorder: Beyond monoamine hypothesis. *Psychiatry and clinical neurosciences* 2018; **72**(1)**:** 3-12.

16.	Culmsee C, Michels S, Scheu S, Arolt V, Dannlowski U, Alferink J. Mitochondria, Microglia, and the Immune System-How Are They Linked in Affective Disorders? *Front Psychiatry* 2018; **9:** 739.

17.	Dean J, Keshavan M. The neurobiology of depression: An integrated view. *Asian journal of psychiatry* 2017; **27:** 101-111.

18.	Liu CH, Zhang GZ, Li B, Li M, Woelfer M, Walter M *et al.* Role of inflammation in depression relapse. *Journal of neuroinflammation* 2019; **16**(1)**:** 90.

19.	Otte C, Gold SM, Penninx BW, Pariante CM, Etkin A, Fava M *et al.* Major depressive disorder. *Nature reviews Disease primers* 2016; **2:** 16065.

20.	Langfelder P, Horvath S. WGCNA: an R package for weighted correlation network analysis. *BMC bioinformatics* 2008; (1)**:** 559-559.

21.	Lachmann A, Rouillard AD, Monteiro CD, Gundersen GW, Jagodnik KM, Jones MR *et al.* Enrichr: a comprehensive gene set enrichment analysis web server 2016 update. *Nucleic Acids Research* 2016; **44**(W1)**:** W90-W97.

**Supplementary Figures**

**Supplementary Figure 1. UCMS-exposed mice show elevated behavioural emotionality.** **A)** Fur quality scores in control (black) and UCMS (red) groups over 5 weeks (n=10/group, all males). Repeated-measures ANOVA revealed a significant effect of time, stress, and stress x time (all p<1x10^-4^). UCMS mice showed a significantly greater coat degradation than controls from week 2 onwards. **B)** UCMS mice show reduced consumption of sucrose (p=2.88x10-7) but no change in water consumption (p=0.233) compared to controls. **C)** UCMS mice showed significantly increased latency to drink a sweet milk solution in a novel environment (p=1.75x10^-6^) compared to controls, and no difference in their home cage (p=0.736). **D)** UCMS mice show a trend-level increase in avoidance of the food zone in the phenotyper box after light challenge compared to controls (p=0.0626). **E-G)** UCMS mice show no differences from controls in the open field, elevated plus maze, or novelty-suppressed feeding tests, respectively. **H)** UCMS mice show no difference in locomotor activity compared to controls (p=0.258). **I)** UCMS mice show significantly elevated behavioural emotionality z-scores (p=7.41x10^-6^), a composite score of all 6 behavioural tests shown in panels B-G, compared to controls. #p<0.1 *p<0.05, **p<0.01, ***p<0.001.

**Supplementary Figure 2. Overall enrichment profiles and analysis of a priori biological functions across cell-types.** **A)** Heatmap of up-regulated (red) and down-regulated (blue) pathways. Summaries above heatmap indicate the total number of up- and down-regulated pathways in each cell-type. **B)** Parent-child association analysis of a priori biological functions. Column “#” denotes the total number of significantly enriched child terms for the respective parent across all cell-types. **C)** Histogram of NES within cell-types, and relative distribution of pathways identified in parent-child analysis. NES of individual pathways are indicated with a tick below x axes.

**Supplementary Figure 3. Density of cortical microcircuit cell types in control and UCMS groups.** Bar graph showing mean cell density of PYR, SST, PV, and VIP-cells in control (black) and UCMS (red) groups. Error bars represent standard error. # indicates a trend-level, non-significant (p=0.090), difference in density of PV-cells between control and UCMS mice.

**Supplementary Tables**

**Supplementary Table 1.** Differential expression results in each cell type. Log2 fold-change, raw p-value, and independent filtering-corrected p-value shown for each cell-type.

**Supplementary Table 2.** Leading-edge analysis of EnrichmentMap-derived pathway clusters in PYR-cells. Differential expression results of genes overlapping across 5 or more pathways within each EnrichmentMap-derived cluster are shown. BaseMean represents the average normalized-counts across control and UCMS groups, padj represents adjusted p-value.

**Supplementary Table 3.** Leading-edge analysis of EnrichmentMap-derived pathway clusters in SST-cells. Differential expression results of genes overlapping across 5 or more pathways within each EnrichmentMap-derived cluster are shown. BaseMean represents the average normalized-counts across control and UCMS groups, padj represents adjusted p-value.

**Supplementary Table 4.** Leading-edge analysis of EnrichmentMap-derived pathway clusters in PV-cells. Differential expression results of genes overlapping across 5 or more pathways within each EnrichmentMap-derived cluster are shown. BaseMean represents the average normalized-counts across control and UCMS groups, padj represents adjusted p-value.

**Supplementary Table 5.** Leading-edge analysis of EnrichmentMap-derived pathway clusters in VIP-cells. Differential expression results of genes overlapping across 5 or more pathways within each EnrichmentMap-derived cluster are shown. BaseMean represents the average normalized-counts across control and UCMS groups, padj represents adjusted p-value.

**Supplementary Table 6.** Enrichr analysis of WGCNA modules correlated with emotionality scores in control and UCMS groups. Enrichment analysis was performed separately for up-regulated and down-regulated genes within each module.
