## Supplementary figures and images for "Chronic stress induces co-ordinated cortical microcircuit cell type transcriptomic changes consistent with altered information processing"

### Supplementary Figure 1

#### Supplementary Figure 1

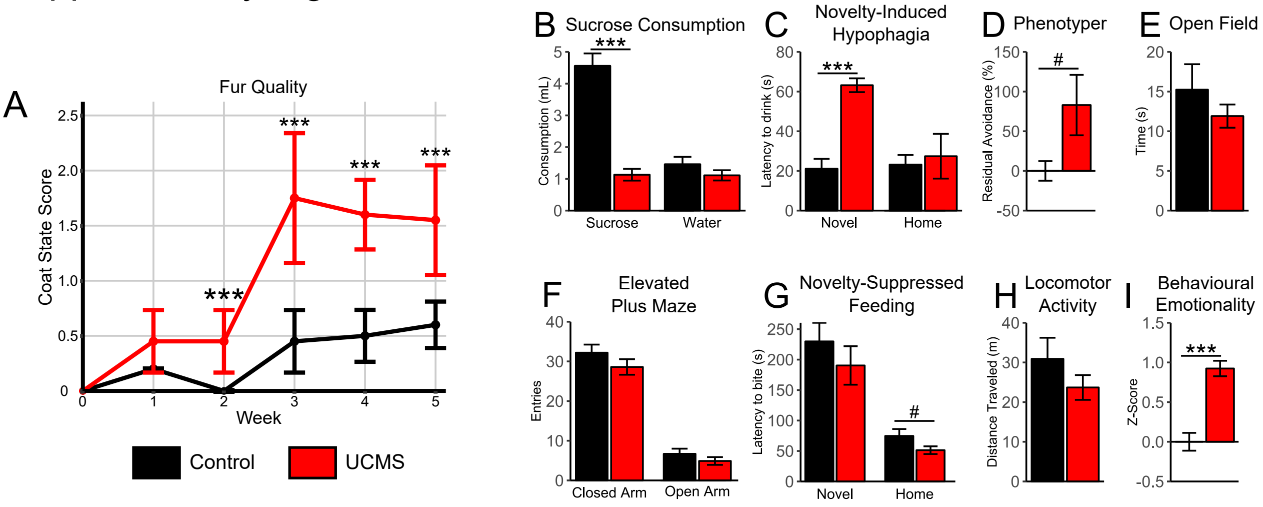

### Supplementary Figure 3

#### Supplementary Figure 3

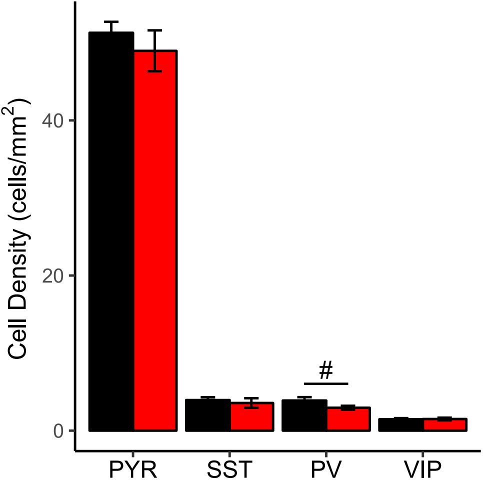
