## Supplementary Figure 2 for "Chronic stress induces co-ordinated cortical microcircuit cell type transcriptomic changes consistent with altered information processing"

A

| Category | GO Term | # | PYR Up | PYR Down | SST Up | SST Down | PV Up | PV Down | VIP Up | VIP Down |
|---|---|---|---|---|---|---|---|---|---|---|
| **Synaptic Transmission** | chemical synaptic transmission | 6 | | 6 | | | | 2 | | |
| | regulation of neuronal signal transduction | | | | | | | | | |
| | postsynaptic neurotransmitter receptor activity | | | | | | | | | |
| | channel regulator activity | 3 | 3 | | | | | | | |
| | signaling receptor activity | | | | | | | | | |
| | neurotransmitter receptor activity | | | | | | | | | |
| | neurotransmitter transporter activity | | | | | | | | | |
| | ion transmembrane transporter activity | 8 | | 1 | | | | 5 | | 2 |
| | neuron differentiation | 20 | 2 | 2 | 2 | | | 5 | | 9 |
| **Synapse Structure** | postsynapse | 2 | | | | | | 1 | | 2 |
| | presynapse | 4 | | 1 | 1 | | | 1 | | 1 |
| | synapse | 11 | | 2 | 1 | 1 | | 2 | | 6 |
| | neuron part | 10 | | 1 | 2 | | | 3 | | 4 |
| | synaptic membrane | 2 | | | | | | 1 | | 2 |
| | synapse organization | 3 | 2 | | | 1 | | | | |
| | synapse assembly | | | | | | | | | |
| | cell adhesion | 11 | 3 | | 9 | | | | | |
| **Inflammation** | immune response | 3 | | 2 | | 1 | | 1 | | |
| | inflammatory response to antigenic stimulus | | | | | | | | | |
| | regulation of immune response | 3 | | 2 | | 1 | | 1 | | |
| | cytokine activity | | | | | | | | | |
| | cytokine receptor activity | | | | | | | | | |
| | inflammatory response | | | | | | | | | |
| **Cellular and Oxidative Stress** | cellular response to stress | 37 | 4 | 6 | 4 | 1 | 4 | 1 | 13 | 4 |
| | cellular response to nitrosative stress | | | | | | | | | |
| | cellular response to oxidative stress | 3 | | | 1 | | | | 2 | |
| | regulation of cellular response to stress | 19 | | 4 | 1 | 1 | 2 | | 8 | 3 |
| | oxidoreductase activity | 5 | | 1 | | | 3 | | 1 | |
| | oxidation-reduction process | 6 | | 2 | | | 4 | | 1 | |
| | cell death in response to oxidative stress | 1 | | | 1 | | | | | |
| | response to stress | 52 | 6 | 6 | 6 | 1 | 5 | 5 | 19 | 7 |
| **Monoamines** | serotonin receptor signaling pathway | | | | | | | | | |
| | serotonin receptor activity | | | | | | | | | |
| | catecholamine binding | | | | | | | | | |
| | catecholamine transport | | | | | | | | | |
| | organic hydroxy compound metabolic process | 3 | 1 | | | | 1 | | | 1 |
| | noradrenergic neuron differentiation | | | | | | | | | |
| | response to monoamine | | | | | | | | | |
| **Bioenergetics** | generation of precursor metabolites and energy | 7 | | 3 | | | 4 | | 1 | |
| | cellular carbohydrate metabolic process | 3 | | | 2 | 1 | | | | |
| | mitochondrion | 7 | | 4 | 1 | | 2 | | | |

B

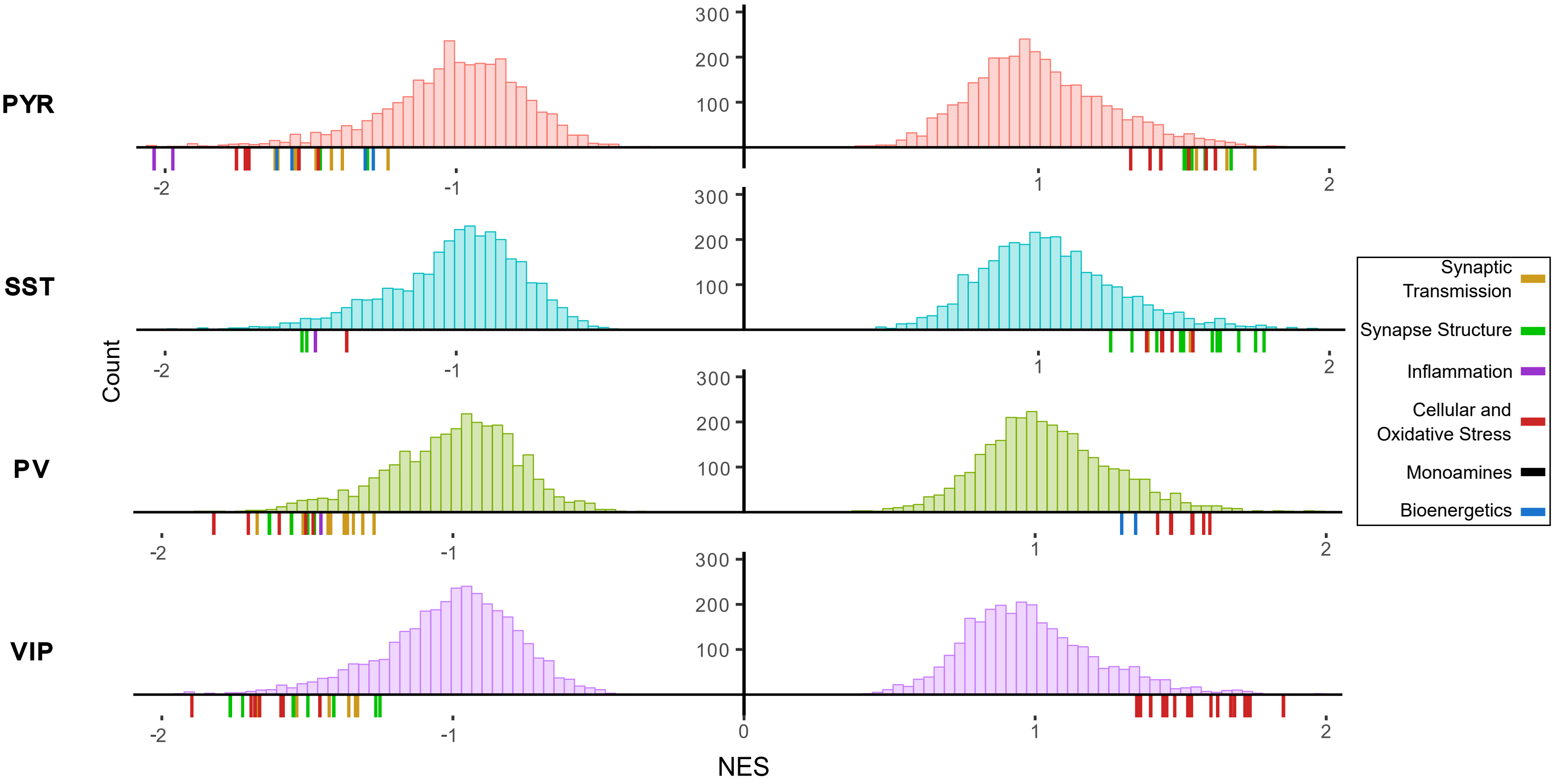
